# Comparative repeat profiling of two closely related conifers (*Larix decidua and Larix kaempferi*) reveals high genome similarity with only few fast-evolving satellite DNAs

**DOI:** 10.1101/2021.03.21.436054

**Authors:** Tony Heitkam, Luise Schulte, Beatrice Weber, Susan Liedtke, Sarah Breitenbach, Anja Kögler, Kristin Morgenstern, Marie Brückner, Ute Tröber, Heino Wolf, Doris Krabel, Thomas Schmidt

## Abstract

In eukaryotic genomes, cycles of repeat expansion and removal lead to large-scale genomic changes and propel organisms forward in evolution. However, in conifers, active repeat removal is thought to be limited, leading to expansions of their genomes, mostly exceeding 10 gigabasepairs. As a result, conifer genomes are largely littered with fragmented and decayed repeats. Here, we aim to investigate how the repeat landscapes of two related conifers have diverged, given the conifers’ accumulative genome evolution mode. For this, we applied low coverage sequencing and read clustering to the genomes of European and Japanese larch, *Larix decidua* (Lamb.) Carrière and *Larix kaempferi* (Mill.), that arose from a common ancestor, but are now geographically isolated. We found that both *Larix* species harbored largely similar repeat landscapes, especially regarding the transposable element content. To pin down possible genomic changes, we focused on the repeat class with the fastest sequence turnover: satellite DNAs (satDNAs). Using comparative bioinformatics, Southern, and fluorescent *in situ* hybridization, we reveal the satDNAs’ organizational patterns, their abundances, and chromosomal locations. Four out of the five identified satDNAs are widespread in the *Larix* genus, with two even present in the more distantly related *Pseudotsuga* and *Abies* genera. Unexpectedly, the EulaSat3 family was restricted to *L. decidua* and absent from *L. kaempferi*, indicating its evolutionarily young age. Taken together, our results exemplify how the accumulative genome evolution of conifers may limit the overall divergence of repeats after speciation, producing only few repeat-induced genomic novelties.

## INTRODUCTION

Ranging in size between 0.002 and nearly 150 Gb, eukaryotic genomes vary by several orders of magnitude (Hidalgo et al., 2017). Among those, conifer genomes are especially large with sizes up to 37 Gb (Ahuja and Neale, 2005). As new reference genome sequences are generated – among them conifers such as spruces, pines, and recently firs and larches – new insights into the composition of conifer genomes are brought forward (Nystedt et al., 2013; Wegrzyn et al., 2014; Stevens et al., 2016; Kuzmin et al., 2019; Mosca et al., 2019). One of the main takeaways is that the steady accumulation of repeats is the main driver for conifer genome expansion, presumably due to limited elimination of transposable elements (TEs; Nystedt et al., 2013; Prunier et al., 2016).

As the large conifer genomes have accumulated repeats over long periods of time with only slow removal and turnover of repetitive sequences, we wondered whether species-specific repeat profiles were able to evolve in closely related conifers. Regarding repetitive sequence classes, it is already hypothesized that TE families likely persist in conifers over long evolutionary timeframes (Zuccolo et al., 2015). In contrast, satellite DNAs (satDNAs) have much faster sequence turnovers than TEs. They form one of the major repeat groups, constituting up to 36 % of some plant genomes (Ambrozová et al., 2011; Garrido-Ramos, 2017). SatDNAs are composed of short monomers with individual lengths often between 160-180 bp and 320-360 bp (Hemleben et al., 2007; Melters et al., 2013), and are arranged in long tandemly repeated arrays. As they confer important functions with roles in cell division, chromatid separation, and chromosome stability (Jagannathan et al., 2018), they often occupy specific chromosomal regions, such as the centromeres and the (sub-) telomeres (Schmidt and Heslop-Harrison, 1998; Melters et al., 2013; Oliveira and Torres, 2018). Due to their fast evolution and defined chromosomal localization, satDNAs may represent valuable targets to trace repeat evolution and divergence over long, evolutionary timeframes in conifers.

As models, we investigate two related conifers within the genus *Larix*, the deciduous European and Japanese larches, i.e. *Larix decidua* (Lamb.) Carrière and *Larix kaempferi* (Mill.). Their genome sizes range between 11 Gb (*L. kaempferi*) and 13 Gb (*L. japonica*) (Zonneveld, 2012; Bennett and Leitch, 2019), with their huge genomes likely being the result of many divergent and ancient repeats (Nystedt et al., 2013; Pellicer et al., 2018). Larches frequently hybridize, leading to an unclear genetic basis with debated phylogenetic positions of individual species (Wei and Wang, 2003; Lu et al., 2014). From a breeding perspective, the interspecific hybrid *Larix* × *eurolepis* (with parental contributions of *L. decidua* and *L. kaempferi*) offers interesting possibilities for larch cultivation outside the natural range, especially in Europe (Pâques et al., 2013), however determining the parental contributions to the traits of larch hybrids remains difficult.

Consistent with other Pinaceae species, all larches have 2n=2x=24 chromosomes with conserved sizes, divided into six meta- and six submetacentric chromosome pairs (Hizume et al., 1993; Prunier et al., 2016). In larches, the 18S-5.8S-25S and 5S rDNAs are physically separated (Garcia and Kovařík, 2013), and fluorescent *in situ* hybridization (FISH) with respective rDNA probes clearly mark three and one chromosome pair for *L. decidua* (Lubaretz et al., 1996) and two and one pair for *L. kaempferi* (Liu et al., 2006; Zhang et al., 2010), respectively. Similarly, a single satDNA family is known (“LPD”), marking a heterochromatic chromosomal band on 22 chromosomes in *L. kaempferi* (Hizume et al., 2002). However, how the accumulative genome evolution mode of conifers affects the landscapes of larch repeats after speciation is not understood by far.

To test this, we sequenced L. *decidua* and *L. kaempferi* in low coverage to quantify, classify, and compare the respective repeat fractions. As satDNAs are typically marked by high sequence turnovers, we expect the highest differences for this repeat class. Using comparative bioinformatics, Southern, and fluorescent *in situ* hybridization, we deeply profiled five selected satDNAs, focusing on their abundance, their higher order arrangements, and chromosomal location. Assessment of their genomic distribution over a wider range of gymnosperms may give insight into the evolutionary age of satDNAs, may allow pinpointing how conifer repeat landscapes have diverged after speciation, and may be used to gather information regarding the parentage of larch hybrids.

## MATERIALS AND METHODS

### Plant material and DNA isolation

Needles and seeds of eleven gymnosperm accessions have been obtained from the Forestbotanical Garden Tharandt (Technische Universität Dresden) and the Staatsbetrieb Sachsenforst (Table 1). DNA was isolated from 2 g of homogenized material from frozen needles using the DNeasy Plant Maxi Kit (Qiagen, Hilden, Germany) according to manufacturer’s instructions. To allow for a more efficient elution of conifer DNA, the incubation time during the elution step was increased to 10 minutes. Purified DNA was eluted into water instead of the provided AE buffer.

**Table 1:** Plant material

| # | Species | Origin<sup>a</sup> | Accession | Plant family |
| --- | --- | --- | --- | --- |
| 1 | *Larix decidua* Mill. | T | 50°58'58.0"N 13°23'44.4"E | Pinaceae |
| 2 | *Larix kaempferi* (Lamb.) Carrière | G | #1041 |  |
| 3 | *Larix gmelinii* (Rupr.) Kuzen. | G | #868 |  |
| 4 | *Larix sibirica* Ledeb. | G | #1340 (5/18) |  |
| 5 | *Pseudotsuga menziesii* var. *viridis* (Schwer.) Franco | T | #1014 Indiv. 2 |  |
| 6 | *Pinus sylvestris* L. | T | [U/1] 504 55 |  |
| 7 | *Picea abies* (L.) H. Karst. | T | [*Pf 1935/35] 217/12 |  |
| 8 | *Abies sibirica* Ledeb. | T | [U/1] 403 a210 Indiv. left |  |
| 9 | *Taxus baccata* L. | T | [U/7] 401 c19 | Taxaceae |
| 10 | *Juniperus communis* L. | T | [*2000/117] 837/1684 | Cupressaceae |
| 11 | *Ginkgo biloba* | T | N 50°58'53.5";E 13°34'27.2" | Ginkgoaceae |

<sup>a</sup>Origin of accession, either from Forest Park Tharandt (T) or Staatsbetrieb Sachsenforst Graupa (G)

For cytogenetics, we have used primary root tips from seeds of *L. decidua* (obtained as selected material for propagation from Staatsdarre Flöha, Partie number 1846, ELA/83704) and *L. kaempferi* seeds (obtained from Niedersächsische Landesforsten, provenance number 83901), as well as roottips from *L*. × *eurolepis* plantlets (clone 56.012.15) obtained from somatic embryogenic cultures from Madlen Walter and Kurt Zoglauer from the Humboldt Universität zu Berlin.

### Sequencing, read clustering, repeat classification, and characterization

For sequencing, we used *L. decidua* and *L. kaempferi* as reference, with accessions as indicated in Table 1. Whole genome sequence libraries with 350 to 500 bp fragment sizes have been generated by Macrogen Inc., followed by Illumina paired end sequencing on Illumina HiSeq2000 and HiSeq2500 machines. The reads were trimmed to the same length (101 bp) using *trimmomatic* (Bolger et al., 2014). We pre-treated and interlaced the read sequences using the custom scripts accompanying the local *RepeatExplorer* installation (*paired_fastq_filtering*.*R* and *fasta_interlacer*, followed by *seqclust*). The reads were quality-trimmed to include only sequences with a Phred score ≥ 10 over 95 % of the read length. Overlapping paired ends have been excluded. We randomly selected five million paired reads from each library and subjected those to comparative clustering with the *RepeatExplorer* software (Novák et al., 2010; Novák et al., 2013) and *TAREAN* (Novák et al., 2017). The resulting clusters were classified by similarity searches against the *Conserved Domain Database* for the functional annotation of proteins (Marchler-Bauer et al., 2011), *RepBase* Update (Jurka et al., 2005), the *REXdb* database (Neumann et al., 2019), and a custom library containing ribosomal, telomeric and plastid sequences. Clusters connected by paired reads and sharing a common annotation have been manually combined to superclusters. Graphic representations as bar and pie charts have been produced with *R* using the *ggplot2* library (Wickham, 2016).

Clusters with a satellite-typical star-like and circular graphical representation (Novák et al., 2010) have been selected for further analysis. Putative monomers were manually detected on the *RepeatExplorer*-derived contigs as well as with the software *Tandem Repeats Finder* (Benson, 1999). Using the putative circular monomers as a template, we iteratively aligned the paired reads against these template sequences to derive more robust consensus sequences (Data S1). Repeat sequences have been compared and characterized using multiple sequence alignments and dotplots of monomers with the packages *MAFFT* (Katoh and Standley, 2013) and *FlexiDot* (Seibt et al., 2018). General sequence investigation was performed with the multi-purpose software *Geneious*6.1.8 (Kearse et al., 2012).

### Repeat quantification by comparative read mapping

To determine the relative abundance of the selected *Larix* repeat families in other gymnosperms, we complemented our own data (*L. decidua* and *L. kaempferi*) with publicly available whole genome shotgun Illumina reads from twelve gymnosperms. From the Pinaceae, these data sets include other larches (*Larix sibirica, Larix gmelinii*), pines (*Pinus taeda, P. sylvestris*, and *P. lambertiana*), spruces (*Picea abies, P. glauca*, and *P. sitchensis*), fir (*Abies sibirica*), and Douglas fir (*Pseudotsuga menziesii*). We also analyzed the genomes of distantly related gymnosperms, such as yew (*Taxus baccata*) and common juniper (*Juniperus communis*). The publicly available reads were obtained from NCBI under the accession numbers SRR8555411, SRR2027118, SRR1054646, ERR268439, SRR2027090, ERR268355, SRR1259615, SRR3100750, ERR268418, ERR268427, and ERR268423. We randomly extracted three million paired reads, and iteratively mapped them against the circular satDNA consensus until it remained unchanged. This alignment to the consensus was performed with the *Geneious*6.1.8 mapping tools (using *medium sensitivity* parameters, Kearse et al., 2012). We graphically represented mapping counts as bubble chart with *R* and *ggplot2* (Wickham, 2016).

### Repeat detection in genome assemblies

If we detected EulaSat1 to EulaSat5 presence in additional genomes, we downloaded the corresponding genome assemblies, if available. These included assemblies of *Pseudotsuga menziesii* (Psme v.1.0 from treegenesdb.org, Neale et al., 2017) and *Abies alba* (Abal v.1.1 from treegenesdb.org, Mosca et al., 2019).Using a local *BLAST* search (Altschul et al., 1990), we have retrieved the five scaffolds with the most hits for each of the satDNA consensuses. In order to assess the organization of the satDNA families, we visualized each scaffold as a dotplot. For visualization purposes, we extracted representative 20 kb regions and generated *FlexiDot* dotplots (Seibt et al., 2018) with the parameters *-k 18 -S 4 -c n -p 0 -A 1*.*5 -T 40 -E 16*.

### PCR and cloning

From the monomeric consensus sequences, outward facing primers have been designed from the *L. decidua* reference (Table 2). For the amplification of satDNA probes for Southern hybridization and FISH, PCR was carried out with the specific primer pairs. PCR reactions with 50 ng plasmid template were performed in 50 µl volume containing 10x DreamTaq buffer and 2.5 units of DreamTaq polymerase (Promega). Standard PCR conditions were 94 °C for 5 min, followed by 35 cycles of 94 °C for 1 min, primer-specific annealing temperature for 30 sec, 72 °C for 1 min and a final incubation time at 72 °C for 5 min. The resulting amplicons have been cloned into the pGEM-T vector (Promega), followed by Sanger sequencing. The clones containing inserts most similar to the satDNA consensus have been chosen for hybridization experiments.

**Table 2:**
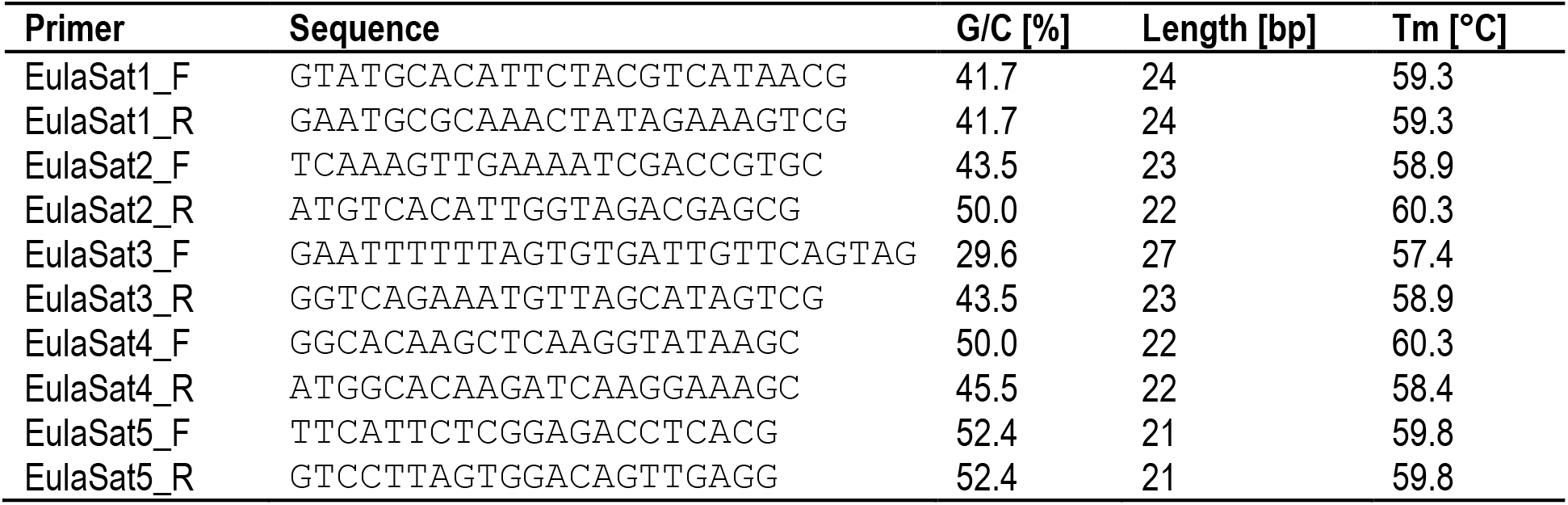
Primer pairs for the generation of satDNA clones

### Southern blot hybridization

For Southern blots, genomic DNA was restricted with enzymes specific for each tandem repeat targeted, separated on 2 % agarose gels and transferred onto Hybond-N+ nylon membranes (GE Healthcare) by alkaline transfer. Hybridizations were performed according to standard protocols using probes labelled with ^32^P by random priming (Sambrook et al., 1989). Filters were hybridized at 60 °C and washed at 60 °C for 10 min in 2x SSC/ 0.1 % SDS. Signals were detected by autoradiography.

### Probe labelling, chromosome preparation, and fluorescent *in situ* hybridization

Sequenced satDNA clones have been used as template for PCR-based labelling with biotin-16-dUTP.The probe pZR18S containing a 5066 bp fragment of the sugar beet 18S-5.8S-25S rRNA gene (HE578879, Paesold et al., 2012) was labeled with DY-415 or DY-647-dUTP (Dyomics) by nick translation. The probe pXV1 (Schmidt et al., 1994) for the 5S rRNA gene was labeled with digoxygenin-11-dUTP by nick translation.

We prepared mitotic chromosomes from the meristems of young primary roots, harvested shortly after germination. Prior to fixation in ethanol:chloroform:glacial acetic acid (2:1:1), root tips were incubated either for 16 h in 2 mM 8-hydroxyquinoline or for 1 hour in nitrous oxide at 10 bar. Fixed plant material was digested for 0.5 to 1.5 hours at 37° C in an enzyme mixture consisting of 2 % (w/v) cellulase from *Aspergillus niger* (Sigma C1184), 4 % (w/v) cellulase Onozuka R10 (Sigma 16419),0.5 % (w/v) pectolyase from *Aspergillus japonicus* (Sigma) P-3026,1 % (w/v) cytohelicase from *Helix pomatia* (Sigma) C-8274, 1% hemicellulase from *Aspergillus niger*(Sigma H2125), and 20 % (v/v) pectinase from *Aspergillus niger* (Sigma P4716) in citrate buffer (4 mM citric acid and 6 mM sodium citrate).The root tips have been washed and transferred to a slide, before maceration with a needle in 45 % glacial acetic acid. Before the slide dried, the chromosomes have been fixed with methanol:glacial acetic acid (3:1).

Before FISH, according to the amount of cytoplasm visible under light microscope, we pre-treated the slides with 100 µg/ml RNase in 2× SSC for 30 min, followed by 200 µl of 10 µg/ml pepsin in 10 mM HCl for 15 to 30 min. Slides with abundant cytoplasm were additionally treated for 10 min with proteinase K. FISH was performed according to the protocol of Heslop-Harrison et al.(1991) with modifications as described (Schmidt et al., 1994). Probes were hybridized with a stringency of 76 % and subsequently washed with a stringency of 79 %. The chromosome preparations were counterstained with DAPI (4’, 6’-diamidino-2-phenylindole) and mounted in antifade solution (CitiFluor). Slides were examined with a fluorescence microscope (Zeiss Axioplan 2 imaging) equipped with Zeiss Filter 09 (FITC), Zeiss Filter 15 (Cy3), Zeiss Filter 26 (Cy5), AHF Filter F36-544 (aqua),and Zeiss Filter 02 (DAPI). Images were acquired directly with the Applied Spectral Imaging v. 3.3 software coupled with the high-resolution CCD camera ASI BV300-20A.

### Data access

Whole genome shotgun Illumina sequences have been deposited at the European Bioinformatics Institute (EBI) short read archive in the project PRJEB42507 (http://www.ebi.ac.uk/ena/data/view/PRJEB42507). Similarly, cloned sequences used as Southern and FISH probes have been deposited at EBI with the accession numbers LR994496 to LR994500).

## RESULTS

### *L. decidua* and *L. kaempferi* show very similar repeat profiles

To assess the genome composition of *L. decidua* and *L. kaempferi*, we obtained paired-end Illumina whole genome shotgun sequences with fragment sizes of 500 bp. Five million reads of each genome have been randomly chosen for comparative low coverage clustering with *RepeatExplorer*. The software automatically chose 2,124,798 (*L. decidua*) and 2,125,214 (*L. kaempferi*) reads (corresponding to a genome coverage between 1.6 and 1.9 %) and yielded estimates of the repetitive fraction of 69.0 % for *L. decidua* and 68.1 % for *L. kaempferi*. We classified the read clusters according to their repeat class and manually combined clusters connected by read pairs and similar annotation to superclusters. This has led to similar repeat profiles for both genomes (Figure 1): In particular, Ty3-*gypsy* retrotransposons (approx. 31 % for both genomes) made up the largest fraction, followed by Ty1-*copia* retrotransposons (both approx. 24 %). Presentation of the first 214 read superclusters as a two-sided, comparative barchart illustrates the high degree of genomic similarity between both genomes (Figure 1). With only few exceptions, the read clusters are equally abundant in *L. decidua* and *L. kaempferi*, with gaps indicating absence of the repeat from one of the genomes. We detected most variation in the amount of satellite DNA, with 3.2 % for *L. decidua* and 2.0 % for *L. kaempferi* (Figure 1, marked in red).

**Figure 1:**
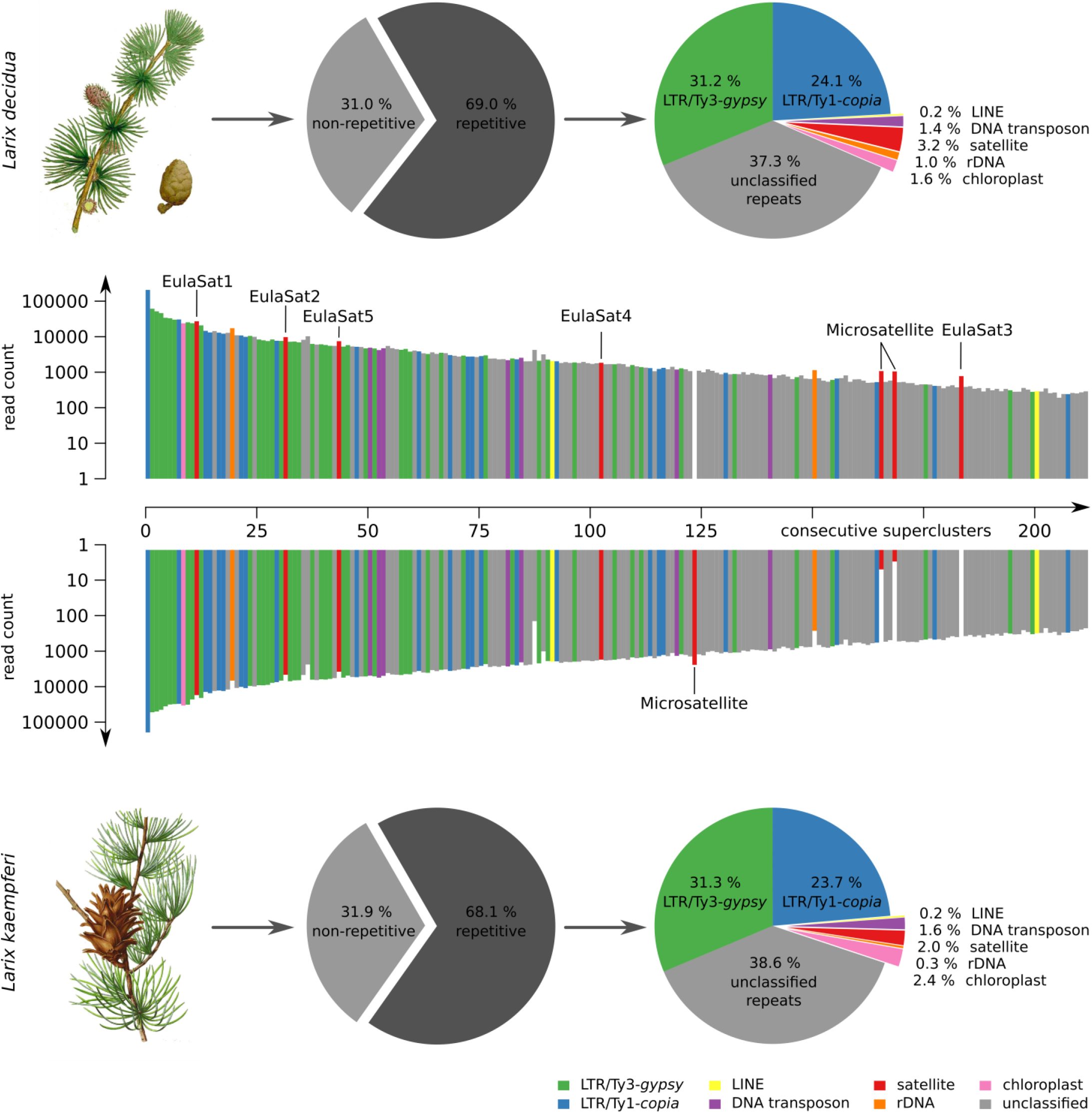
Comparison of repetitive genome fractions reveals high genomic similarity over all repeat types between *L. decidua* and *L. kaempferi*. At the center of each figure, a two-sided barplot shows 214 repeat superclusters with respective read counts in *L. decidua* (top) and *L. kaempferi* (bottom). The read count is presented on a logarithmic scale. Experimentally analyzed clusters are marked with the corresponding repeat family name. The composition of each *Larix* repeat fraction is summarized by pie charts. The plant illustrations are reproduced from Woodville, W., Hooker, W.J., Spratt, G., Medical Botany, 3th edition, vol. 1: (1832) (*L. decidua*) and M. E.-A. Carriere (ed.) Revue Horticole, serié 4, vol. 40: (1868), Paris (*L. kaempferi*).

### *Larix* tandem repeats vary in abundance and genome organization, with only punctual differences between *L. decidua* and *L. kaempferi*

Six of the analyzed *RepeatExplorer* read clusters produced circular or star-shaped layouts, typical for tandem repeats (Figure S1), representative of five satDNA families. Using *L. decidua* as reference organism, we extracted sequences of the candidates, and named them EulaSat1 to EulaSat5, short for <u>Eu</u>ropean <u>la</u>rch <u>sat</u>ellite. We refined the monomer consensus sequences by iterative mapping of three million paired reads to generate robust consensus sequences (Figure 2, Data S1), used for quantification and primer generation. In order to verify the consensus sequence and to generate hybridization probes, we amplified and cloned all five candidates from *L. decidua*. SatDNA characteristics are summarized in Table 3, whereas a multi-sequence dotplot shows the family and subfamily structure (Figure S2).

**Table 3:** Characteristics of satDNA in *L. decidua* (*Ld*) and *L. kaempferi* (*Lk*) genomes.

| Family | Genome proportion [%]<sup>a</sup> |  | Monomer length [bp] |  | G/C content [%] |  | Mean pairwise identity [%] |  | Read support |  |
| --- | --- | --- | --- | --- | --- | --- | --- | --- | --- | --- |
|  | Ld | Lk | Ld | Lk | Ld | Lk | Ld | Lk | Ld | Lk |
| EulaSat1 | 1.28 | 0.81 | 173 | 173 | 33 | 33 | 91.8 | 90.3 | 74,373 | 46,173 |
| EulaSat2 | 0.46 | 0.22 | 148 | 148 | 45 | 45 | 84.7 | 83.9 | 22,881 | 10,714 |
| EulaSat3a | 0.05 | 0.00 | 345 | 345 | 28 | 28 | 98.0 | --- | 1,532 | 0 |
| EulaSat3b |  |  | 345 | 345 | 31 | 31 | 94.2 | --- | 978 | 0 |
| EulaSat3c |  |  | 345 | 345 | 26 | 26 | 95.2 | --- | 605 | 0 |
| EulaSat4a | 0.09 | 0.08 | 203 | 203 | 43 | 43 | 86.6 | 85.8 | 1,104 | 696 |
| EulaSat4b |  |  | 169 | 169 | 43 | 43 | 86.8 | 86.3 | 264 | 119 |
| EulaSat5 | 0.35 | 0.18 | 87 | 87 | 50 | 50 | 87.1 | 84.8 | 4,510 | 1,398 |

<sup>a</sup> RepeatExplorer-based estimate
<sup>b</sup> Number of reads mapping out of three million paired end reads

**Figure 2:**
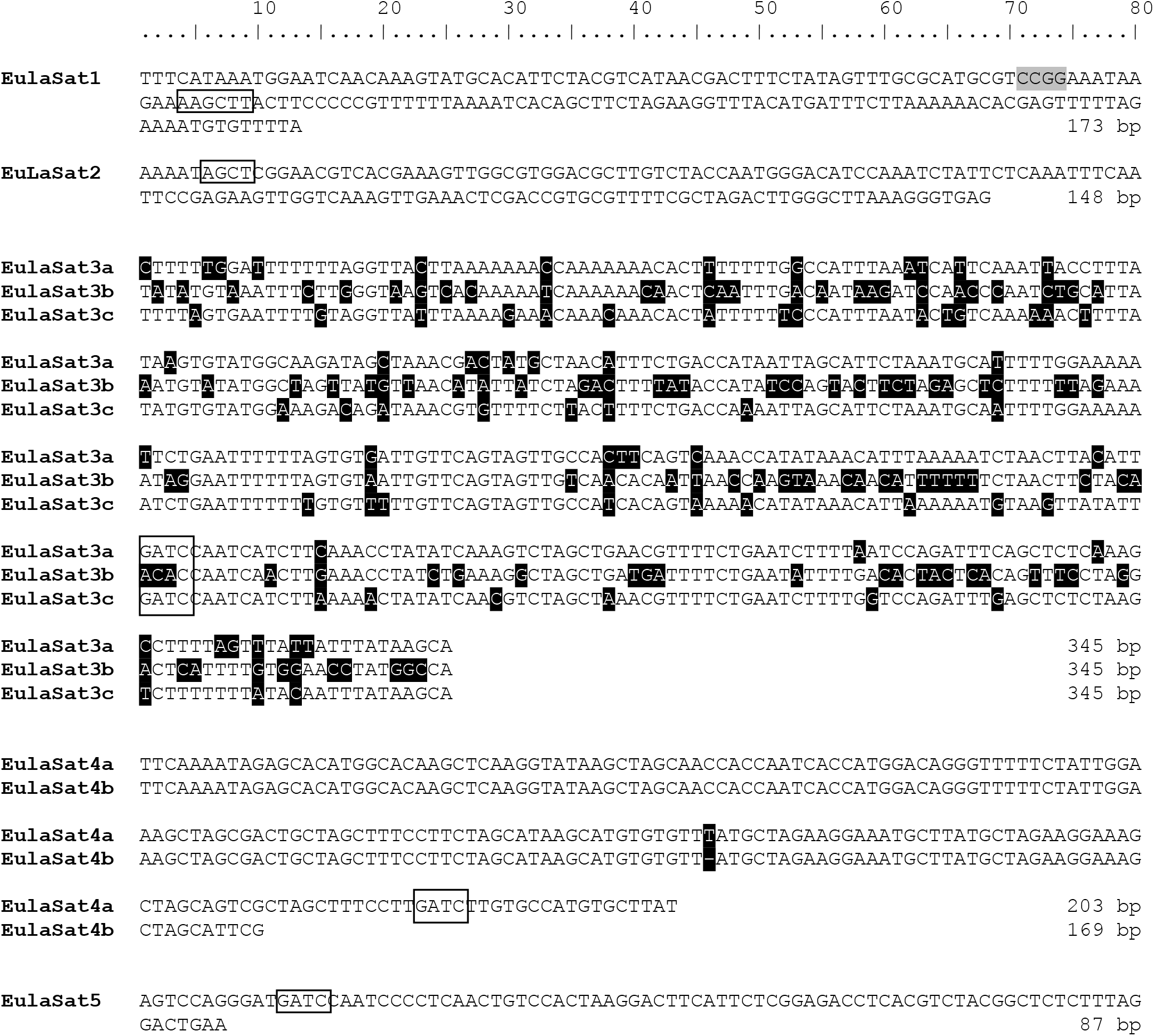
Consensus sequences and subunit structure of the tandem repeat monomers. The monomer consensus sequences of the EulaSat1 to EulaSat5 satDNAs are shown. Recognition sites of restriction enzymes used to release the DNA ladder (Figure 3, Figure 5) are indicated by rectangles. *Hpa*I/*Msp*I recognition sites are shaded in grey. EulaSat3 and EulaSat4 are divided into the EulaSat3a, EulaSat3b and EulaSat3c as well as the EulaSat4a, and EulaSat4b subfamilies. These sequences are represented as multiple sequence alignment, with ambiguities shaded in black.

To verify the head-to-tail organization of the five EulaSat repeat families, we transferred restricted genomic *L. decidua* DNA onto Southern membranes. After Southern hybridization of the EulaSat probes, we investigated the resulting autoradiographs for presence of satDNA-typical ladder hybridization (Figure 3), indicating repeat organization in long arrays. In detail, summarizing the computational and molecular data, the five satDNAs are characterized as follows:

**Figure 3:**
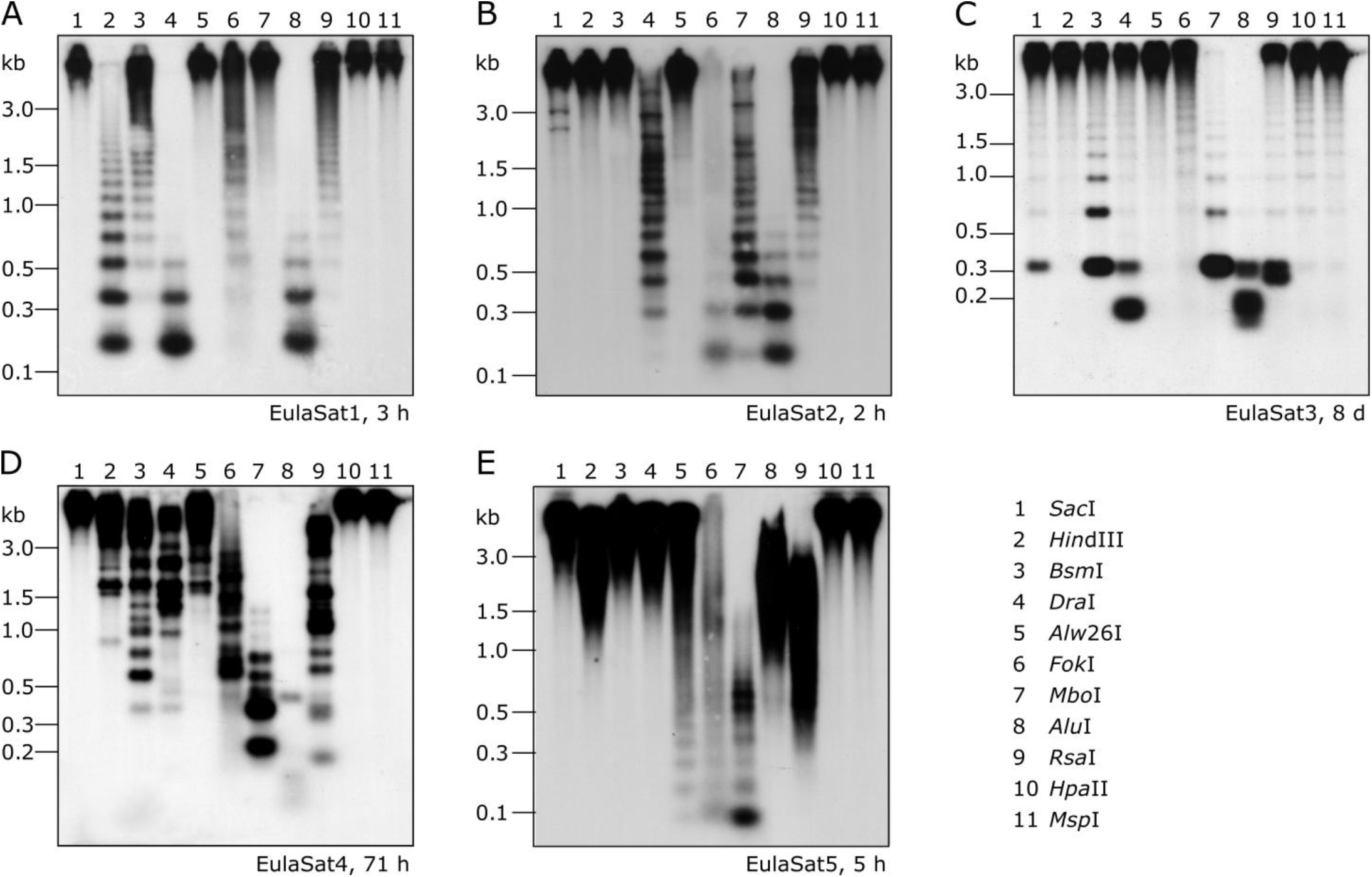
Genomic organization of the *Larix* satDNA families EulaSat1 to EulaSat5. Southern hybridization of restricted genomic *L. decidua* DNA with satDNA probes releases the satellite-typical ladder pattern of EulaSat1 **(A)**, EulaSat2 **(B)**, and EulaSat3 **(C)**, EulaSat4 **(D)**, and EulaSat5 **(E)**. The exposition time are indicated in hours (h) or days (d) for each experiment.

1. Comprising approx. 1 % of both *Larix* genomes, EulaSat1 is the major satDNA family in larches. First described as LPD, it is an integral part of many *Larix* genomes (Hizume et al., 2002), forming conserved 173 bp monomer with a G/C content of 33 %. Six of the eleven enzymes tested produced a satellite-typical restriction ladder for EulaSat1, all supporting the monomer length of 173 bp (Figure 3A). Clearest signals up to the tetra- and pentamers have been generated with *Dra*I (lane 4) and *Alu*I (lane 8), whereas the remaining enzymes such as *Hin*dIII (lane 2) released longer multimers up to the dodecamer. Although the EulaSat1 consensus contains a potential *Hpa*II/*Msp*I restriction site (indicated in Figure 2), we detected only high molecular weight signals, indicating a high degree of DNA methylation.
2. With a genomic representation of 0.46 % and 0.22 %, EulaSat2 is the second-most abundant satDNA family in *L. decidua* and *L. kaempferi*. EulaSat2 has a relatively high G/C content with 44 % and consists of monomers with the satDNA-typical length of 148 bp. The EulaSat2 autoradiograph (Figure 3B) showed ladder-like patterns for five enzymes. *Fok*I, *Mbo*I, and *Alu*I (lanes 6-8) released the EulaSat2 monomer, supporting its length of 148 bp. *Dra*I (lane 4) only produced weak monomeric signals, whereas *Rsa*I (lane 9) did not generate any monomeric bands, pointing to only weak restriction site conservation. Bands up to the undecamer were released, before falling together in a smear. Hybridization of *Hpa*II/*Msp*I restricted DNA did not produce any signals below 3 kb.
3. The EulaSat3 family is divided into three subfamilies with similar features: The conserved 345 bp long monomers contain a generally low G/C content between 26 % and 31 %. Out of all identified repeats, only the three EulaSat3 subfamilies are genome-specific, as their clusters contain only reads from *L. decidua* and none from *L. kaempferi*. Consensus sequences of all EulaSat3 subfamilies can be subdivided into a 178 bp and a 167 bp subunit with identities ranging between 45.5 % and 48.3 % (Figure S3), suggesting evolution by EulaSat3 reorganization into structures of higher order. EulaSat3 hybridization (Figure 3C) generated ladder-like patterns with different intensities in all lanes, with its monomeric length (345 bp) distinguishable in most cases. For two enzymes, *Dra*I (lane 4) and *Alu*I (lane 8), bands below the monomer size were visible. These additional bands can be explained by multiple restriction sites in the monomer (see Figure 2), giving rise to 163 and 182 bp fragments (*Dra*I) as well as 36, 176, 196 and 212 bp fragments (*Alu*I). In addition, *Hpa*II and *Msp*I were able to cut EulaSat3, both producing identical, weak ladders (lanes 10-11), pointing to presence of at least some monomers without DNA methylation in the putative restriction site.
4. Similarly, for EulaSat4, we detected two subfamilies with different monomeric lengths. EulaSat4a has 203 bp monomers and is more abundant, supported by a mapping of 1,104 reads. In contrast, the less frequent EulaSat4b subfamily (supported by 696 reads) has a monomer length of 169 bp. We did not detect clear, canonical ladder patterns after hybridization of EulaSat4 (Figure 3D). However, signals as detected for *Bsm*I (lane 3), *Mbo*I (lane 7), and *RsaI* (lane 9) can be explained by the recognition of both EulaSat4 subfamilies by the Southern probe. As observed, a combination of 203 and 169 bp fragments leads to the complex ladder patterns with unequal step sizes.
5. Out of all identified satDNA families, EulaSat5 has the shortest monomer (87 bp) and the highest G/C content (50 %). Although the monomer is short, this satDNA family makes up 0.35 % and 0.18 % of the *L. decidua* and the *L. kaempferi* genomes, respectively.EulaSat5 hybridization (Figure 3E) yielded ladder patterns for the three enzymes *AIw*26I, *Fok*I and *Mbo*I (lanes 5-7). For *Mbo*I, a strong monomeric signal was detected, providing additional support for the monomer size of 87 bp and for the high restriction site conservation within EulaSat5 arrays. Intense signals in the hexa- and heptamer regions indicate arrays with higher order repeat structures. Hybridization of *Hpa*II/*Msp*I-restricted DNA did not reveal bands in the low molecular weight region, suggesting strong EulaSat5 DNA methylation.

### Individual *Larix decidua* chromosomes show comparable satDNA localizations

To determine the position of the satDNA families along *Larix* chromosomes, we prepared mitotic and interphase chromosomes from the *L. decidua* reference, and *in situ* hybridized them with biotin-labeled satDNA probes (Figure 4A-E):

**Figure 4:**
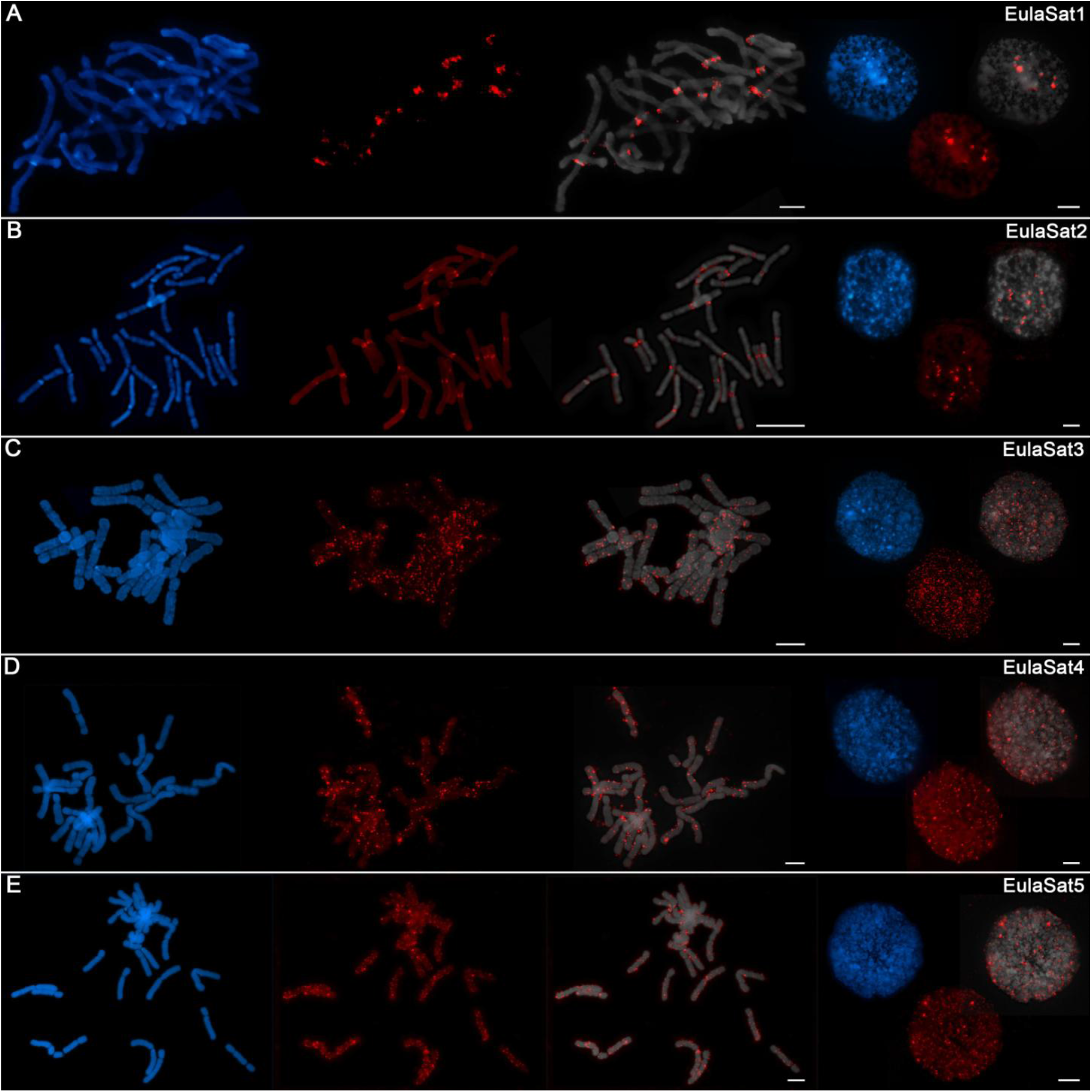
Chromosomal localization of the EulaSat families along *Larix decidua* chromosomes. Chromosomes have been counterstained with DAPI, indicated in blue and grey. Fluorescent *in situ* hybridizations of EulaSat1 (A), EulaSat2 (B), EulaSat3 (C), EulaSat4 (D), and EulaSat5 (E) to *L. decidua* meta- and interphases are shown in red.

1. EulaSat1 hybridized to 18 from the 24 *L. decidua* chromosomes, co-localizing with the strongly DAPI-stained heterochromatic proximal bands (Figure 4A). EulaSat1’s occurrence in the deep heterochromatin was confirmed by co-localization with DAPI-positive regions on interphase nuclei (Figure 4A).
2. For EulaSat2, we have observed presence on all chromosomes. The localization along the centromeric constriction of all chromosomes indicates EulaSat2’s suitability to serve as a marker for the centromere (Figure 4B). As this position is depleted in DAPI staining, we assume that the EulaSat2 regions are only loosely packaged. At higher resolution, using interphase nuclei, we confirmed that EulaSat2 is largely excluded from the heterochromatin (Figure 4B).
3. The three remaining satDNA families, EulaSat3 to EulaSat5 are marked by a dispersed localization along all *L. decidua* chromosomes (Figure 4C-E). For EulaSat3, we identified a range of minor signals without exclusion of the centromeres. Nevertheless, we found an enrichment of EulaSat3 hybridization sites along the intercalary and distal chromosome regions. At interphases, we noted EulaSat3 presence in hetero- and euchromatic regions (Figure 4C).
4. This pattern is mirrored for EulaSat4. We found that most of the minor EulaSat4 signals were localized at the intercalary chromosome regions. The distal chromosome regions and the centromeric restrictions were not excluded, but only few chromosomes carried EulaSat4 signals at these regions. At interphases, most signals were localized in the DAPI-positive heterochromatin (Figure 4D).
5. EulaSat5 signals were scattered over the whole length of all chromosomes, with frequent enrichments at or near the (peri-)centromeric regions. The signals are often euchromatic, but without exclusion from the DAPI-positive heterochromatin (Figure 4E).

Taken together, whereas three of the satDNA probes (EulaSat3 to EulaSat5) are dispersed along all chromosomes, EulaSat1 and EulaSat2 produce distinct signals, limited to the heterochromatic band and the centromeric constriction, and produce clear chromosomal landmarks.

### Distribution, abundance, and genomic organization in related conifer genomes

We used bioinformatics and experimental approaches to investigate the abundance and genomic organization of the EulaSat repeats in related species. Using a read mapping approach, we screened whole genome shotgun Illumina reads of twelve Pinaceae species (Figure 5), including four larches, three pines, three spruces, a fir, and a Douglas fir. As outgroups, we also analyzed DNA of more distantly related yew (*Taxus baccata*) and juniper (*Juniperus communis*) trees.

**Figure 5:**
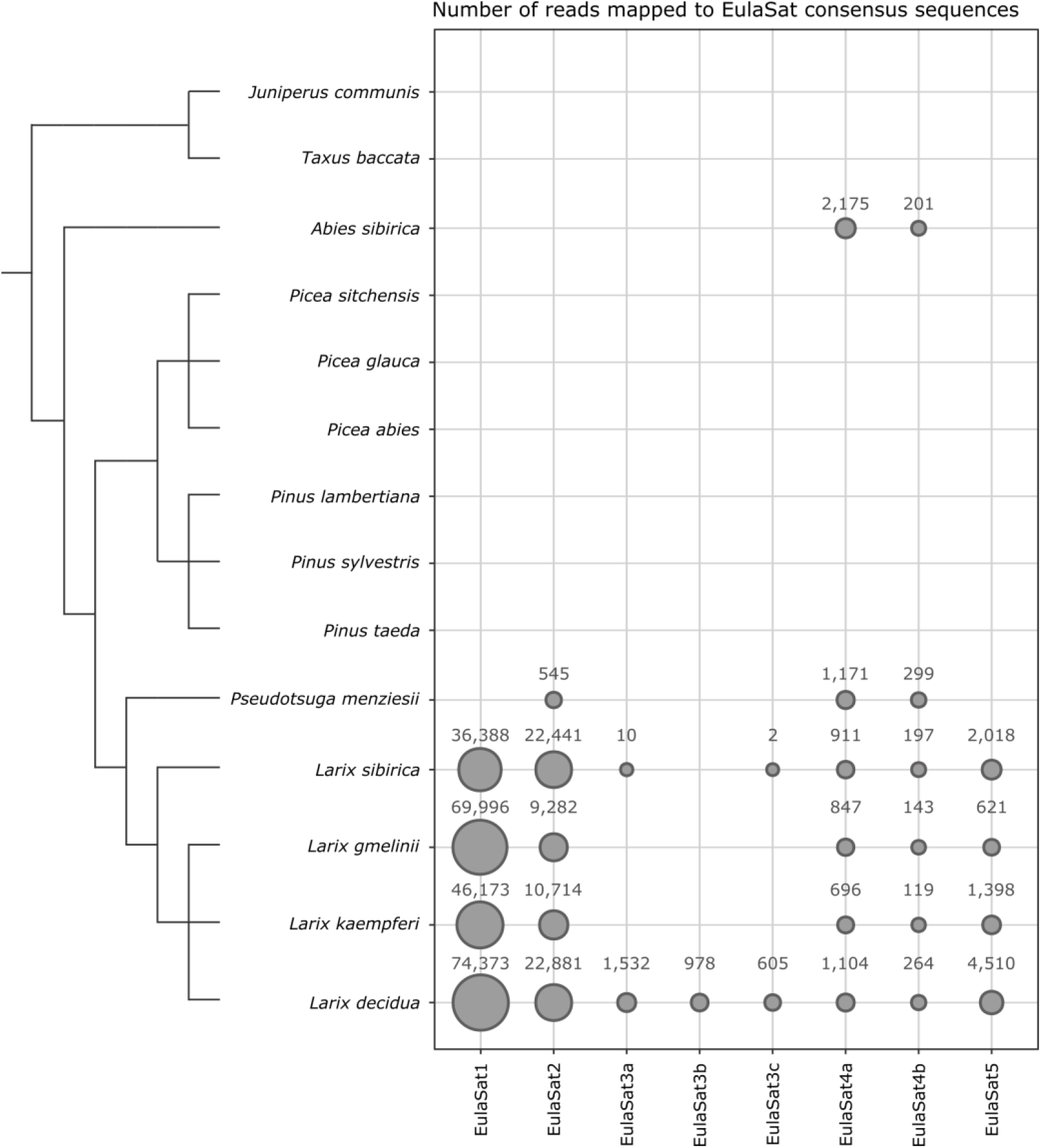
Distribution of the EulaSat tandem repeats in fourteen gymnosperm genomes surveyed by read mapping. The area of each bubble represents the amount of whole genome shotgun Illumina reads mapping to the EulaSat consensus sequences. A total of three million paired reads has been used as input for the mapping analysis. The dendrogram indicates the evolutionary relationship between the species according to Wei et al. (2003) for the genus *Larix* and Lu et al. (2014) for the overall phylogeny. The branch lengths are not to scale.

As read mapping may misrepresent the factual genome representation of repeats due to inherent G/C biases (Benjamini and Speed, 2012; Chen et al., 2013), we complemented our bioinformatics approach with an experimental verification. For this, we comparatively hybridized the satDNA probes onto restricted genomic DNA, and quantified the repeat abundance in eleven species (Figure 6). Our species sampling includes *L. decidua, L. kaempferi, L. gmelinii, L. sibirica* (lanes 1-4), and a single representative of additional gymnosperm genera: *Pseudotsuga menziesii* (lane 5), *Pinus sylvestris* (lane 6), *Picea abies* (lane 7), *Abies sibirica* (lane 8), *Taxus baccata* (lane 9), *Juniperus communis* (lane 10), and *Ginkgo biloba* (lane 11).

**Figure 6:**
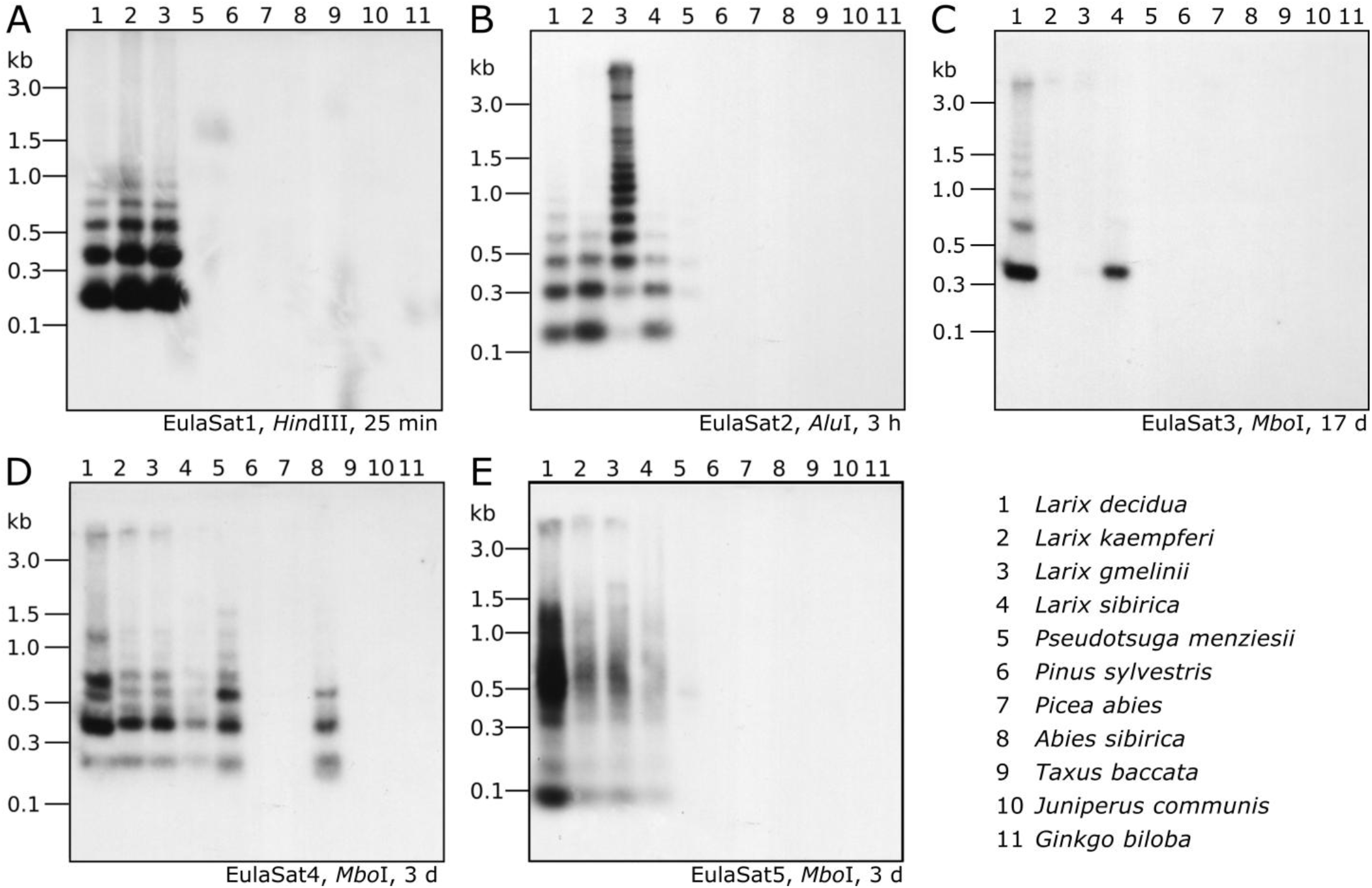
Organization and abundance of EulaSat repeats in related gymnosperm genomes. Genomic DNA of eleven gymnosperms has been restricted as indicated in each panel and was analyzed by comparative Southern hybridization of EulaSat1 **(A)**, EulaSat2 **(B)**, EulaSat3 **(C)**, EulaSat4 **(D)**, and EulaSat5 **(E)**. Exposure times ranged between 25 minutes and seventeen days, as indicated below the autoradiographs.

Both approaches show that EulaSat1, EulaSat2, EulaSat4, andEulaSat5 are present in all four *Larix* accessions analyzed, indicating their wide-spread occurrence throughout the genus (Figures 5, 6).

1. Especially EulaSat1 is highly abundant in all four *Larix* species, but without occurrence outside of the genus (Figure 5). Supporting this, EulaSat1 hybridization revealed clear ladder signals in the genus *Larix*, already after 25 min of exposition (Figure 6A). We observed similar patterns and signal strengths in all *Larix* species tested, indicating similar EulaSat1 monomer sizes with organization in long arrays across the genus. The remaining genomes did not produce any signal, pointing to EulaSat1 absence. Longer exposition time of 3 hours revealed no further information.
2. A similar high abundance in *Larix* sp. was detected for EulaSat2. EulaSat2 was also present in *P. menziesii*, but in lower quantity (Figure 5). After EulaSat2 hybridization, clear ladder-like pattern is visible for all larch species tested (Figure 6B), supporting the organization of similar-sized monomers in a tandem arrangement. In addition, for *Pseudotsuga menziesii* (lane 5), very weak signals corresponding to the dimer and trimer are distinguishable, becoming more prominent after longer exposure (not shown), without additional signals in any other lanes. Hybridization to *L. gmelinii* DNA (lane 3) does only produce faint monomeric and dimeric bands, and instead leads to many signals in the higher, multimeric region. As the *L. gmelinii* DNA was restricted completely, this indicates a less conserved *Alu*I restriction site in the EulaSat2 satDNA.
3. Computationally, the three EulaSat3 subfamilies have been analyzed individually, indicating considerable genomic impact only in *L. decidua*. We did not detect presence in *L. kaempferi* and *L. gmelinii*. For subfamilies EulaSat3a and EulaSat3c, only few *L. sibirica* hits mapped to the consensus, suggesting a reduced abundance in this genome. The other gymnosperm sequences tested did not contain any similarity to the EulaSat3 subfamilies (Figure 5). The patchy distribution across the *Larix* genus was also apparent experimentally (Figure 6C), with hybridization revealing exclusive signals in *L. decidua* and *L. sibirica*. In both species, the monomeric band constituted the strongest signal, suggesting the high conservation of the *Mbo*I restriction site within EulaSat3. In *L. decidua* the satDNA-typical ladder pattern was formed, whereas in *L. sibirica* the multimeric bands were absent. As the signals were still faint after 17 days of exposure, we conclude a relatively low abundance in both genomes.
4. Out of all satDNAs analyzed, the EulaSat4a and EulaSat4b subfamilies had the broadest distribution. Apart from their presence in the *Larix* genomes, they also populate *P. menziesii* and *A. alba* genomes. In all six EulaSat4-containing genomes EulaSat4a has been more abundant than EulaSat4b (Figure 5). Corroborating this, the corresponding autoradiograph showed signals in species of the *Larix, Pseudotsuga*, and *Abies* genera (Figure 6D, lanes 1-5, 8). The remaining Pinaceae species (*Pinus sylvestris* and *Picea abies*) did not carry any signals, with longer exposition time (seven days) not changing this result. Hybridization to the larches produced very similar patterns, pointing to similar genomic organization. In *Abies sibirica* (lane 8) the lowest band represents a double signal, presumably generated by conserved *Mbo*I restriction sites in the two EulaSat4 subrepeats (169 and 203 bp). However, hybridization to *P. menziesii* (lane 5) produced a stronger ladder with bands slightly shifted towards lower molecular weights, suggesting a small deletion within the EulaSat4 monomers in this species.
5. Read mappings indicate EulaSat5 restriction to *Larix* genomes, with highest abundance in *L. decidua* (Figure 5). However, the corresponding probe hybridized to the species of the *Larix* and the *Pseudotsuga* genera (Figure 6E, lanes 1-5). Signal patterns of the larch species tested resemble each other, with a relatively strong monomeric band, a fainter dimeric band, and a smear at a higher molecular weight. In *P. menziesii* (lane 5), the smear was overlaid by a very faint band at approximately 480 bp, indicating low abundance. Longer exposition (six days) of the autoradiograph did not reveal EulaSat5 in further species.

Experimental and computational approaches revealed that two satDNA families also occurred outside of the *Larix* genus, i.e. in *Pseudotsuga* (EulaSat2, EulaSat4) and *Abies* (EulaSat4). For both genera, genome assemblies were available (Neale et al., 2017; Mosca et al., 2019). We queried those with all satDNA consensuses, and deeply inspected the five scaffolds with the most satDNA hits in the genome assemblies of *P. menziesii* and *A. alba*:

In *P. menziesii*, we extracted long EulaSat2 arrays spanning scaffolds over a megabase, with and without higher order arrangements (Figure S4A). This indicates that the EulaSat2 family, though less abundant (Figure 5, 6B) still plays a major role in this genome.

For EulaSat4, in *P. menziesii*, we detected some arrays over 20 kb, often interrupted by other repeats (Figure S4B). The arrays included variable monomers and different homogenization with or without higher order. In *A. alba*, longer arrays have been detected more frequently. Strikingly, we noticed less monomer variation, with stronger homogenization and a higher abundance of EulaSat4a than EulaSat4b (Figure S4C).

Taken together, our three approaches (read mapping, analyses of genome assemblies, and experimental quantification) corroborate the different abundances of the five satDNA repeats in the gymnosperms. We confirmed presence of EulaSat1, EulaSat2, EulaSat4, and EulaSat5 in all *Larix* genomes tested. EulaSat2 and EulaSat4 reside also in more distantly related Pinaceae genomes. In contrast, experimental and bioinformatic evidence support the young age of EulaSat3 that is restricted to Siberian and European larches.

### Only very few differences distinguish the chromosomes of *L. decidua* from those of *L. kaempferi*

We aimed to combine the information gained from *in situ* hybridization to *L. decidua* chromosomes (Figure 4) as well as from the quantitative comparisons of conifer genomes (Figures 5-6). We now asked, how *L. decidua* and *L. kaempferi* genomes differ on a chromosomal scale and if this information can be used to determine the parentage of individual chromosomes in hybrids.

Therefore, we have comparatively hybridized the most promising tandem repeat landmark probes onto metaphases of both larch species (Figure7A-D), including also the 5S and 18S-5.8S-25S rDNA probes and the satDNAs EulaSat1 and EulaSat2.

To check how the rDNA tandem repeat loci compare, we investigated the localization of the 5S and 18S-5.8S-25S rDNAs (Figure 7A-B). Both species harbor two 5S rDNA sites (magenta), located distally at the chromosome arms. For the 18S-5.8S-25S rDNA, we observed hybridization on three chromosome pairs for *L. decidua*, and two pairs for *L. kaempferi* (green), all localized at the secondary constrictions of the chromosomes (Figure 7A-B).

**Figure 7:**
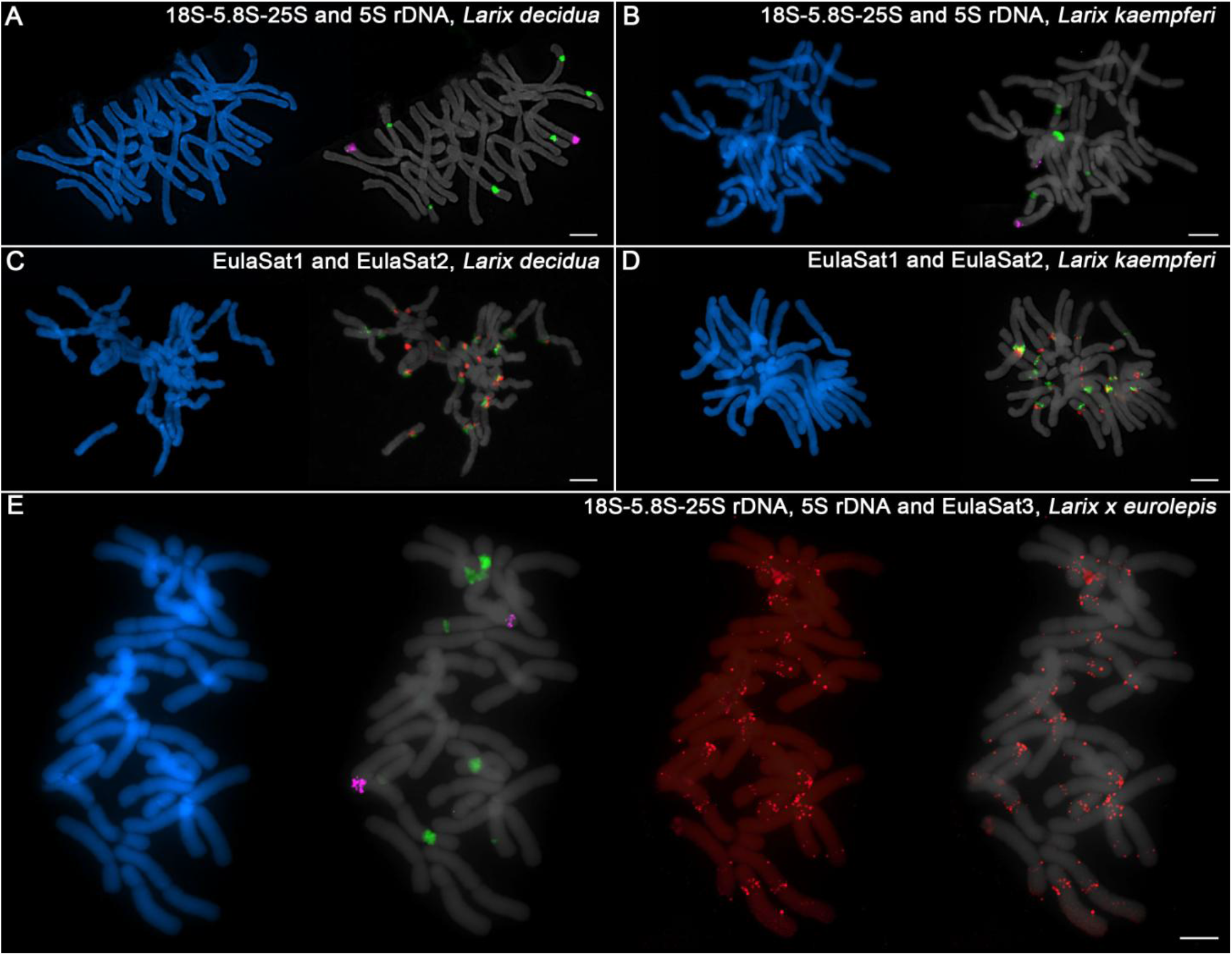
Chromosomal location of rDNAs and the EulaSat families for comparison of *L. kaempferi, L. decidua*, and *L. × eurolepis*. The chromosomes have been counterstained with DAPI, indicated in blue and grey. Reproduced are fluorescent *in situ* hybridizations of the 5S (magenta) and 18S-5.8S-25S rDNAs (green) to metaphases of *L. decidua* (A) and *L. kaempferi* (B). EulaSat1 (green) and EulaSat2 (red) were comparatively hybridized along metaphase chromosomes of *L. decidua* (C) and *L. kaempferi* (D). The genome-specific EulaSat3 family (red) was hybridized along chromosomes of the interspecific *L. × eurolepis* hybrid, along with probes for the 5S (magenta) and 18S-5.8S-25S rDNAs (green).

Regarding the EulaSat1 and EulaSat2 satDNA families, a comparative hybridization onto *L. decidua* and *L. kaempferi* metaphases showed that the satDNA arrays bordered for both species, but with limited co-localization (Figure 7C-D). Overall, the comparison between the major satDNAs EulaSat1 and EulaSat2 yielded only very few differences between both species.

We then shifted attention to the genome-specific, but dispersed EulaSat3 satDNA family that may be used to discern the parentage of individual chromosomes in hybrids. For this, we have prepared metaphases from *Larix* × *eurolepis*, a hybrid between *L. decidua* and *L. kaempferi* (Figure 7E). Hybridization of the 5S (magenta) and 18S-5.8S-25S rDNAs (green) have yielded two and five signals, respectively, with the uneven 18S-5.8S-25S rDNA site number being a testimony to the hybrid status of the individual. The EulaSat3 hybridization yields chromosomes with dispersed EulaSat3 hybridization, indicating *L. decidua* heritage, as well as chromosomes without signals, pointing to descendance from *L. kaempferi*. Nevertheless, due to the dispersed pattern, theEulaSat3 satellite can only give clear parental information for few chromosomes and should be complemented by additional markers, if any become available.

Summarizing, genomes and chromosomes of European and Japanese larches are very similar, with only very few hallmark differences. These include the number in rDNA sites and the genome-specific satDNA family EulaSat3.

## DISCUSSION

### Similar repeat profiles in European and Japanese larch genomes presumably result from repeat accumulation versus turnover

Large conifer genomes evolve only slowly and keep many of their genomic repeats buried within the genomes. With only limited downsizing, we hypothesized that two closely related conifer genomes (such as those from European and Japanese larches) may not accumulate many changes in their overall repeat landscapes. To test this, we have investigated the repeat profiles of these related larch genomes, starting with a broad repeat comparison and then focusing on the repeat class with the fastest sequence turnover, the satDNAs.

We have applied short read sequencing followed by read clustering to efficiently gain insights into both genomes’ satDNA contents (as laid out by Weiss-Schneeweiss et al., 2015; Novák et al., 2017). This approach has been successfully used to characterize the repeat landscapes of many non-model plant species as for example beans, various grasses, camellias, crocuses, quinoa, and ferns (Cai et al., 2014; Heitkam et al., 2015; Ávila Robledillo et al., 2018; Kirov et al., 2018; Liu et al., 2019; Schmidt et al., 2019; Heitkam et al., 2020; Ribeiro et al., 2020), and also of non-model animals such as locusts, grasshoppers, or fishes (Ruiz-Ruano et al., 2016; Ferretti et al., 2020; Boštjančić et al., 2021). For larch genomes, we provided evidence that LTR retrotransposons and derived fragments are their main components, well in line with reports for the related pines and spruces (Kamm et al., 1996; Kossack and Kinlaw, 1999; Nystedt et al., 2013; Stevens et al., 2016; Perera et al., 2018).

As only highly repetitive sequences ≥ 90 % are considered in the *RepeatExplorer* cluster analysis, the size estimations of *Larix* repeat fractions (approximately 68 % of the analyzed genomes) are bound to be vast underrepresentations, excluding the more fragmented repeats. Especially in large genomes, such as those of the conifers analyzed here, fragmentation and slow repeat divergence lead to barely recognizable transposable elements (TEs), often termed “dark matter” (Maumus and Quesneville, 2016). With increasing genome sizes, these dark matter repeats accumulate, leading to the observed and potentially misleading low repeat fraction estimates, as also recently highlighted by Novák et al. (2020).

For *L. decidua* and *L. kaempferi*, we find overall strikingly similar repeat profiles, especially regarding the TE content, without major differences between European and Japanese larches. The similarities include both, repeat family and abundance. In line with evidence from other conifers (Prunier et al., 2016), these results also suggest the limited TE elimination, usually carried out by recombination, reshuffling, or removal as aftermath to genomic rearrangements (Ma et al., 2004; Ren et al., 2018; Kögler et al., 2020; Maiwald et al., 2020). Along the same lines, we did not observe any transpositional bursts of amplification during the speciation of the larches. Thus, only limited TE-induced genomic novelty has likely occurred in the larches’ accumulative genome landscapes.

### Evolutionarily young and old satDNAs contribute to genomic novelty in giant larch genomes

We asked whether the conifer’s genomic background of transposable element accumulation and fragmentation has impacted the evolution of satDNAs. These usually evolve by continued rounds of mutation and fixation leading to relatively fast sequence turnovers, even at structurally important chromosomal locations such as the centromeric regions (Dawe, 2005; Plohl et al., 2012). Indeed, the satDNA abundances in the analyzed larch genomes differed much more than the respective TE portions: We estimated the total satDNA content of *L. decidua* to 3.2 % and of *L. kaempferi* to 2.0 %, corresponding to satDNA amounts of approximately 416 and 220 Mb, respectively. Overall, the larch satDNA proportions are of the same order of magnitude as already estimated from BACs and fosmids for the related pines (1 %; Wegrzyn et al., 2013; Neale and Wheeler, 2019). However, compared to the large values the satDNA genome fraction can occupy in angiosperms (up to 36 %; Ambrozová et al., 2011), the relative amount for larches is rather low, though also not uncommon for angiosperm genomes (Garrido-Ramos, 2017).

To better understand the contribution of satDNA to the genomic differences in larches, we investigated five satDNA families in detail. Here, we will discuss their evolutionary trajectories ranging from the evolutionarily oldest families occurring in several conifer genera (EulaSat2 and EulaSat4), over to those distributed only in the genus *Larix* (EulaSat1 and EulaSat5), to the species-specific family EulaSat3:

The most widely distributed satDNA identified is EulaSat4, with occurrences in larches, Douglas fir and Siberian fir. Interestingly, the comparative read mappings, Southern hybridizations and analyses of available genome assemblies point to longer and more homogenized EulaSat4 arrays in common and Douglas firs than those observed in larches. We therefore think that EulaSat4 is an evolutionarily old repeat, probably playing a larger role in the *Abies* and *Pseudotsuga* genomes. EulaSat4’s patchy distribution across the conifers is an example of a satDNA family’s occurrence that is incongruent with the species phylogeny. The satellite library hypothesis may explain this pattern, by assuming that a common set of satDNAs resides in genomes in low copy numbers (Fry and Salser, 1977; Utsunomia et al., 2017; Palacios-Gimenez et al., 2020). Different satDNA amplification would then lead to the observed patchy abundance pattern of EulaSat4.These low-copy satDNAs may reside within transposable elements, possibly using these for their conservation and amplification (McGurk and Barbash, 2018; Belyayev et al., 2020; Vondrak et al., 2020). EulaSat4’s dispersed localization along all *L. decidua* chromosomes as well as the complex *RepeatExplorer* cluster graphs may indicate such a retrotransposon association. As retrotransposons are strongly conserved across conifer species (Zuccolo et al., 2015; this report), it is likely that EulaSat4 has been retained within a transposable element, followed by patchy amplification in *Larix, Pseudotsuga*, and *Abies* species.

The EulaSat2 family co-localizes with the primary constriction of the *L. decidua* and *L. kaempferi* chromosomes, indicating a possible role in centromere formation. Although some plants have centromeres that differ fundamentally from each other (Gong et al., 2012), that does not seem to be the case for larches. The centromeres of all chromosomes harbor EulaSat2, indicating similar sequences and structures. The 148 bp monomers of EulaSat2 are well in line with lengths observed for other centromeric satDNAs, such as that of rice (Zhang et al., 2013), and a bit shorter than the canonical ∼170 bp monomers of the mammalian alpha satellite (Willard and Waye, 1987). EulaSat2 is more abundant in *L. decidua* than in *L. kaempferi*, indicating recent array size fluctuations. Nevertheless, EulaSat2 is evolutionarily older with presence in the related Douglas fir, but absence from the more distantly related pine, spruce and fir species tested. In fact, centromeric satDNAs of related spruces have already been characterized and differ strongly from Eulasat2 in sequence and monomer length (305 bp; Sarri et al., 2008; Sarri et al., 2011).

In all larches analyzed, the most abundant satDNA is EulaSat1, also known as LPD (Hizume et al., 2002). Its canonical monomer length of 173 bp is similar in all *Larix* species analyzed, but was not detected outside the genus. It is generally assumed that the most abundant satDNA localizes at the centromeres (Melters et al., 2013), however some exceptions have been already reported, e.g. for camellias (Heitkam et al., 2015). Instead of the expected centromeric locations, EulaSat1 constitutes the highly heterochromatic, DAPI-positive band present on most of its 24 chromosomes. Regarding EulaSat1’s evolution, our data indicates strong EulaSat1 amplification after the split from *Pseudotsuga*. Interestingly, the different species set tested by Hizume et al. (2002) indicates also a patchy abundance in some *Picea, Pinus, Abies*, and *Tsuga* species – claims that we cannot verify with our data. Nevertheless, we can convincingly show that differences in abundance between *L. decidua* and *L. kaempferi* point to EulaSat1 array expansions and reductions during the more recent evolutionary events.

In contrast to EulaSat1 and EulaSat2, only short arrays were detected for EulaSat5, the second satDNA family restricted to the larches. *In situ* hybridization marked a scattered localization along all chromosomes, typical for short satDNA arrays. As with EulaSat4, an explanation for the short arrays may be an association with transposable elements. We have observed partially mixed *RepeatExplorer* clusters that may point towards an embedment within retrotransposons, also often observed for short satDNA arrays (Meštrović et al., 2015; Satović et al., 2016; Belyayev et al., 2020; Sultana et al., 2020). Similarly, concatenated TEs or part from TEs may have satDNA-like properties, but tend to occur dispersedly along chromosomes (Maiwald et al., 2020; Vondrak et al., 2020). Here, our data are not sufficient to conclusively resolve the large-scale organization of EulaSat5 within larch genomes.

In contrast to all other families investigated, EulaSat3 has experienced a very recent birth, indicating an evolutionarily young age. EulaSat3 is clearly absent from *L. kaempferi*, but occurs in *L. decidua* with three subfamilies, all containing distinct 345 bp monomers. Their arrangement in higher order is detectable by close inspection of the monomer consensuses and the autoradiograms after Southern hybridization, indicating still ongoing homogenization. FISH and hybridization to *Hpa*II/*Msp*I-restricted genomic DNA have indicated that at least some EulaSat3 monomers are embedded in euchromatic regions. We speculate that these genomic regions are still actively restructured and recombined; processes that potentially restrict the EulaSat3 array size. Taken together, EulaSat3 is a relatively young satellite DNA and presumably marks the evolutionarily young regions of *L. decidua* genomes.

To investigate whether EulaSat3 can be applied as a chromosome-specific marker of *L. decidua* parentage in hybrid offspring, we have tested, if chromosome regions from *L. decidua* can be identified in *L. × eurolepis* hybrids between *L. kaempferi* and *L. decidua*. Although the *in situ* hybridization clearly marks some chromosome regions as derived from *L. decidua*, this method is not as useful as hoped for the clear differentiation of parentally derived regions along larch hybrid chromosomes.

Nevertheless, EulaSat3’s genome specificity within the *Larix* genus as well as the differences in abundance for many of the remaining satDNAs indicates that even large, highly repetitive genomes with slow sequence turnovers can yield new, evolutionarily young repeats and generate sequence innovation to further genome evolution. If these repeats also carry a phylogenetic signal and may be used for taxonomic means (e.g. as suggested by Dodsworth et al., 2014) is still open.

## Conclusion

As conifers largely accumulate transposable elements with only reduced active removal processes, their genomes become huge, loaded with many fragmented, barely recognizable repeat copies. As a result, we believe that closely related conifers harbor very similar repeat landscapes. We have tested this hypothesis for two larch species and detected highly similar TE profiles as well as very few differences in their tandem repeat compositions. Nevertheless, despite the high overall repeat similarity, we detected EulaSat3, a satDNA family present in European larches, but absent from their Japanese counterparts. This illustrates that repeat-driven genome innovation still plays a role, even in the huge, repetitive and fragmented conifer genomes.

## ACKNOWLEDGEMENTS

We remember the late Thomas Schmidt, who has initiated the comparative analyses of larch genomes and has made the ongoing cooperation between three of the partner institutes listed here possible. We also keep in mind the late Ines Walter who has established cytogenetics for larches in our lab. As a tribute, we have included her very first, serendipitous FISH on larches (Figure 7A).

We gratefully acknowledge funding by the German Federal Ministry of Food and Agriculture (Fachagentur Nachwachsende Rohstoffe e. V.; grants 22031714 and 22002216) to TS, DK, and HW. We thank the Forest Botanical Garden of Tharandt, the Staatsbetrieb Sachsenforst, and Madlen Walther of the Humboldt Universität zu Berlin for the contribution of plant material. We thank Ulrike Herzschuh (Alfred Wegener Institute Potsdam) for providing pre-publication access to sequencing data of *Larix gmelinii*, and we acknowledge Prof. Stefan Wanke for supporting our research initiative.

Furthermore, we acknowledge the TU Dresden Center for Information Services and High Performance Computing (ZIH) for computer time allocations.

## SHORT LEGENDS FOR SUPPORTING INFORMATION

**Figure S1:** Comparative clustering of *L. decidua* and *L. kaempferi* reads yields satDNA-typical cluster graphs, representative for the EulaSat1 to EulaSat5 repeats.

**Figure S2:** All against all dotplots indicating similarity between the satDNA consensus sequences.

**Figure S3**: Higher-order arrangement of EulaSat3 monomers.

**Figure S4:** Dotplots of satDNA-containing scaffolds of *P. menziesii* and *A. alba*.

**Data S1:** Consensus sequences of the EulaSat monomers in fasta format.

## AUTHOR CONTRIBUTIONS

TH and TS coordinated the project and interpreted the results. TH, LS, and BW wrote the manuscript, and all authors contributed, proof-read, and edited. TH and AK coordinated the genome sequencing process. TH performed the bioinformatics analysis and contributed to probe preparation. LS and BW performed Southern experiments, LS, SL, and SB prepared metaphase spreads and performed FISH. KM, MB, UT, HW and DK selected and provided plant material, and guided the project intellectually.

## DISCLOSURE DECLARATION

The authors declare no conflict of interests.

## Notes

https://www.ebi.ac.uk/ena/browser/view/PRJEB42507

### Competing Interest Statement

The authors have declared no competing interest.

