## Supplemental Figures 1-4, Supplemental Data 1 for "Comparative repeat profiling of two closely related conifers (*Larix decidua and Larix kaempferi*) reveals high genome similarity with only few fast-evolving satellite DNAs"

Figures S1 – S4
Data S1

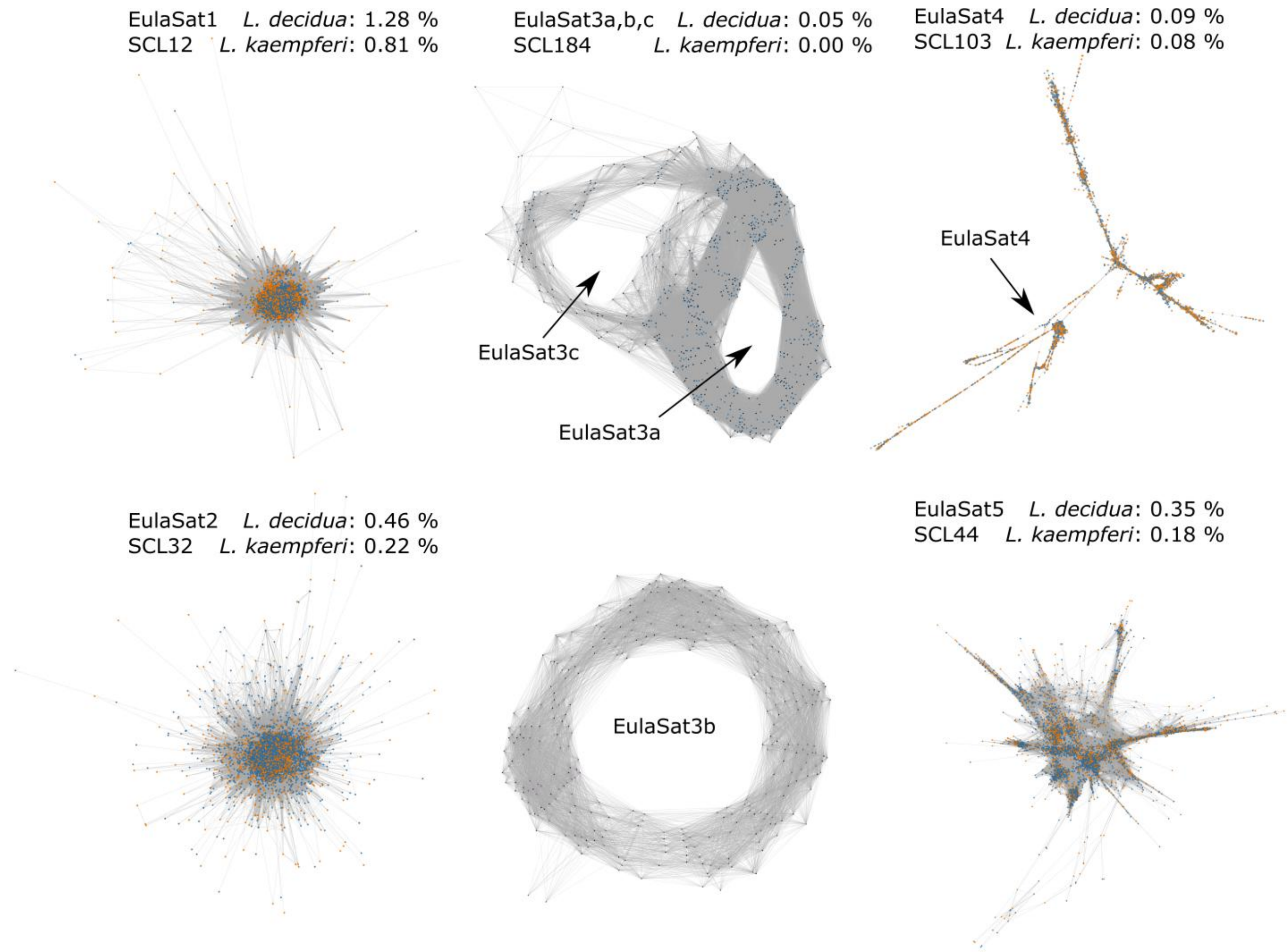


**Figure S1: Comparative clustering of *L. decidua* and *L. kaempferi* reads yields satDNA-typical cluster graphs, representative for the EulaSat1 to EulaSat5 repeats.** Star-like or circular shapes indicate repetitions in a tandem manner. The cluster graphs have been generated by *RepeatExplorer*-based comparative read clustering, with reads from the *L. decidua* and *L. kaempferi* genomes colored as blue and orange nodes, respectively.

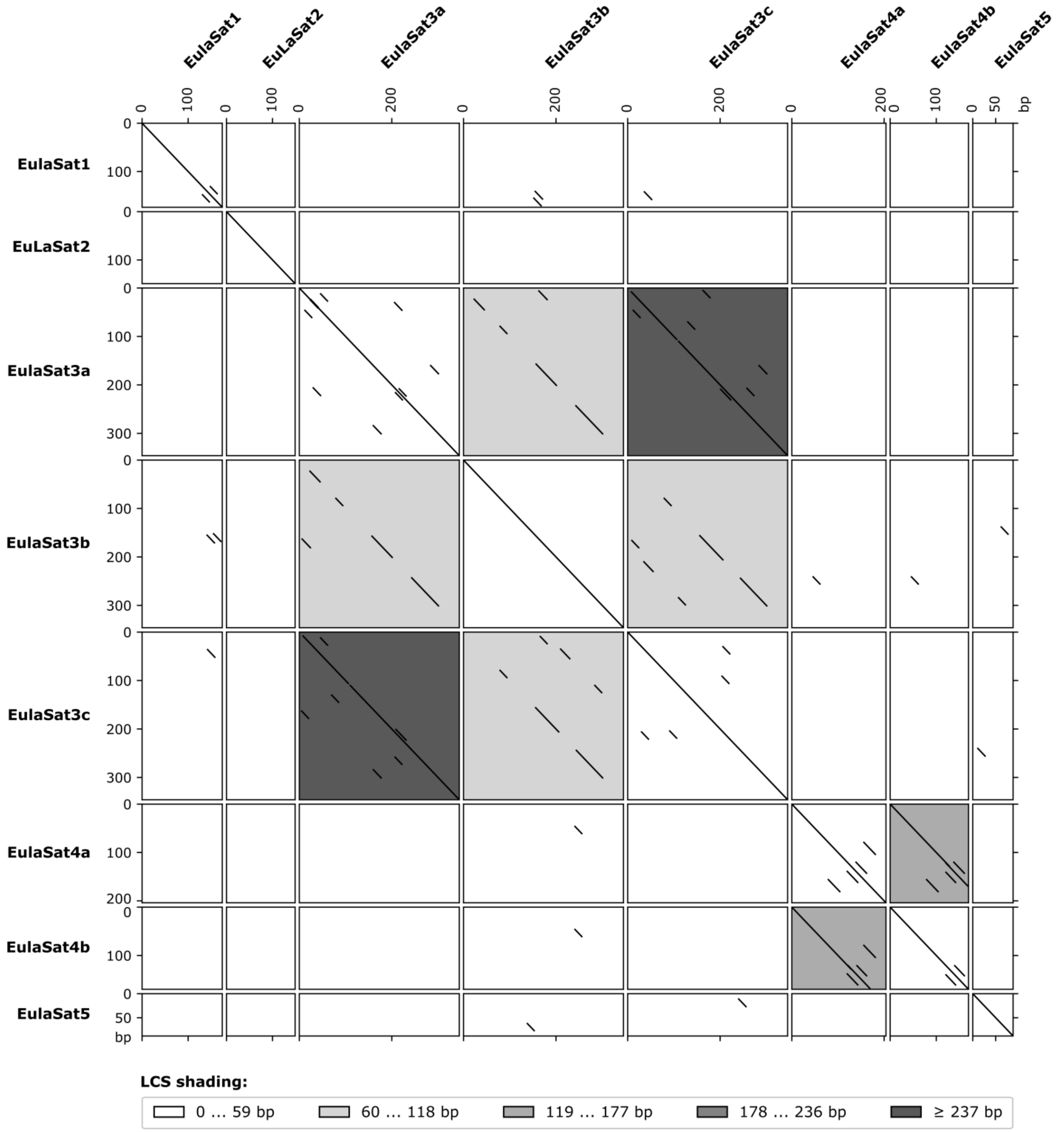


**Figure S2: All against all dotplots indicating similarity between the satDNA consensus sequences.** The figure shows eight self dotplots (main diagonal) and 2×28 pairwise dotplots (below and above the main diagonal). Sequence similarities exceeding 18 bp with allowance of four mismatches are displayed as parallel lines. Pairwise dotplots are shaded according to the length of their longest common subsequence. Here, higher similarity between the subfamilies EulaSat3a, EulaSat3b, and EulaSat3c as well as EulaSat4a and EulaSat4b is visible by shared diagonal lines and dotplot shading.

```
                             10        20        30        40        50        60        70        80        90       100
                     ....|....|....|....|....|....|....|....|....|....|....|....|....|....|....|....|....|....|....|....|
EulaSat3a subunit1   CTTTTTGGATTTTTT----------TAGGTTACTTAAAAAAACCAAAAAAACACTTTTTTTGGCCATTTAAATCATTCAAATTACCTTTATAAGTGTATGG
EulaSat3b subunit1   TATATGTAAATTTCT----------TGGGTAAGTCACAAAAATCAAAAAACAACTCAATTTGACAATAAGATCCAACCCAATCTGCATTAAATGTATATGG
EulaSat3c subunit1   TTTTAGTGAATTTTG----------TAGGTTATTTAAAAGAAACAAACAAACACTATTTTTTCCCATTTAATACTGTCAAAAAACTTTTATATGTGTATGG

EulaSat3a subunit2   TTTTTTAGTGTGATTGTTCAGTAGTTGCCACTTCAGTCAAACCATATAAACATTTAAAAATCTAACTTACATTGATCCAATCATCTTCAAACCTATATCA
EulaSat3b subunit2   TTTTTTAGTGTAATTGTTCAGTAGTTGTCAACACAATTAACCAAGTAAACAACATTTTTTTCTAACTTCTACAACACCAATCAACTTGAAACCTATCTGA
EulaSat3c subunit2   TTTTTTTGTGTTTTTGTTCAGTAGTTGCCATCACAGTAAAAACATATAAACATTAAAAAATGTAAGTTATATTGATCCAATCATCTTAAAAACTATATCA

                            110       120       130       140       150       160       170
                     ....|....|....|....|....|....|....|....|....|....|....|....|....|....|....|...
EulaSat3a subunit1   CAAGATAGCTAAACGACTATGCTAACATTTCTGACCATAATTAGCATTCTAAATGCATTTTTGGAAAAATTCTGA--A
EulaSat3b subunit1   CTAGTTATGTTAACATATTATCTAGACTTTTATACCATATCCAGTACTTCTAGAGCTCTTTTTTAGAAAATAGGA--A
EulaSat3c subunit1   AAAGACAGATAAACGTGTTTTCTTACTTTTCTGACCAAAATTAGCATTCTAAATGCAATTTTGGAAAAAATCTGA--A

EulaSat3a subunit2   AAGTCTAGCTGAACGTTTTCTGAATCTTTTAATCCAGATTTCAGCTCTCAAAGCCTTTTAGTTTATTATTTATAAGCA
EulaSat3b subunit2   AAGGCTAGCTGATGATTTTCTGAATATTTTGACACTACTCACAGTTTCCTAGGACTCATTTTGTGGAACCTATGGCCA
EulaSat3c subunit2   ACGTCTAGCTAAACGTTTTCTGAATCTTTTGGTCCAGATTTGAGCTCTCTAAGTCTTTTTTTATACAATTTATAAGCA
```

**Figure S3**: **Higher-order arrangement of EulaSat3 monomers.** The EulaSat3 monomers can be subdivided into two subunits with a 178 bp + 167 bp arrangement with similarities ranging between 45.5 and 48.3 % to each other. A multiple sequence alignment of the subunits is shown, with ambiguities shaded in black.

**A** EulaSat2 on *Pseudotsuga menziesii* scaffolds (exemplary extractions of 20 kb)

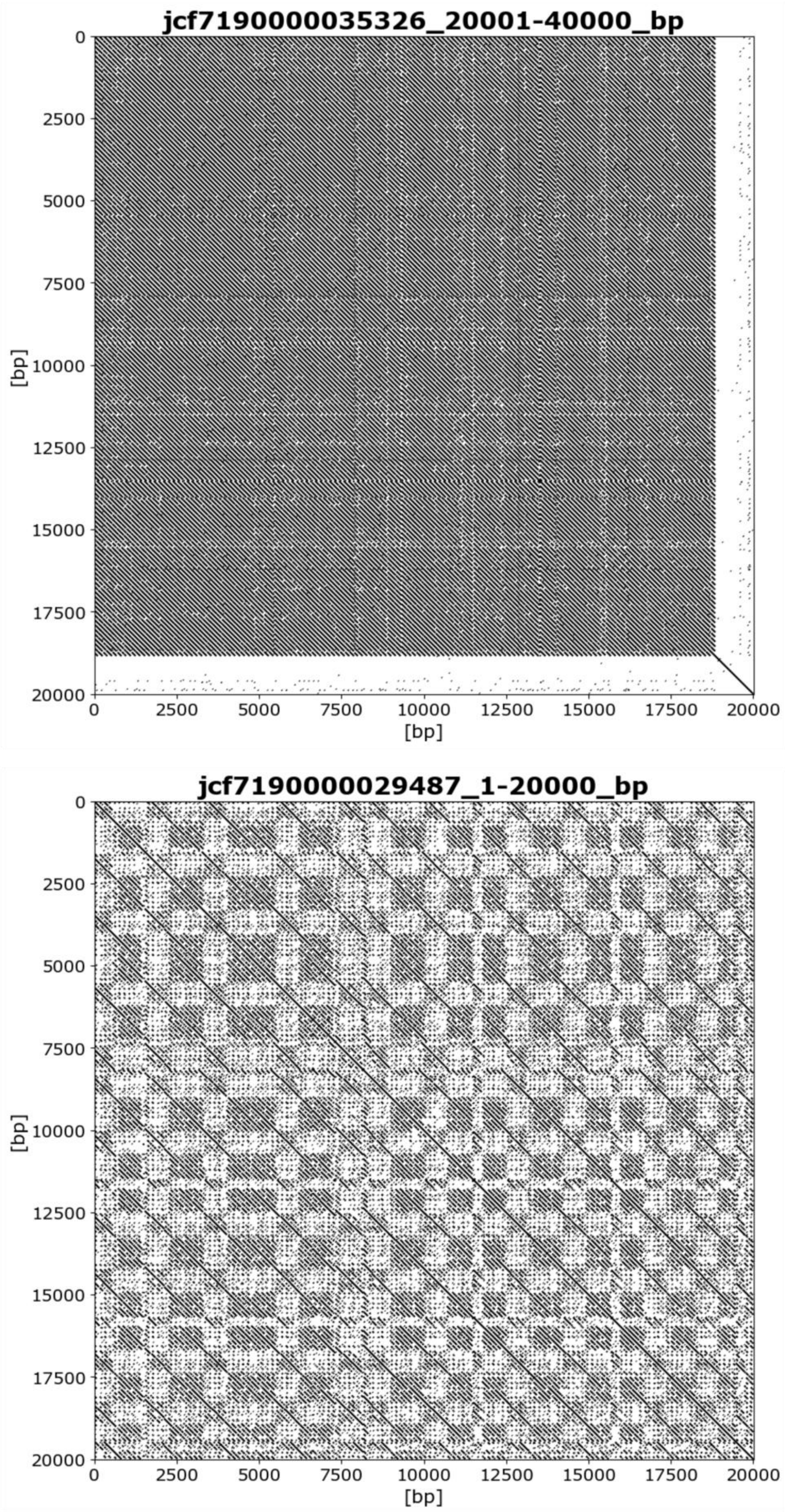

**B** EulaSat4 on *Pseudotsuga menziesii* scaffolds (exemplary extractions of 20 kb)

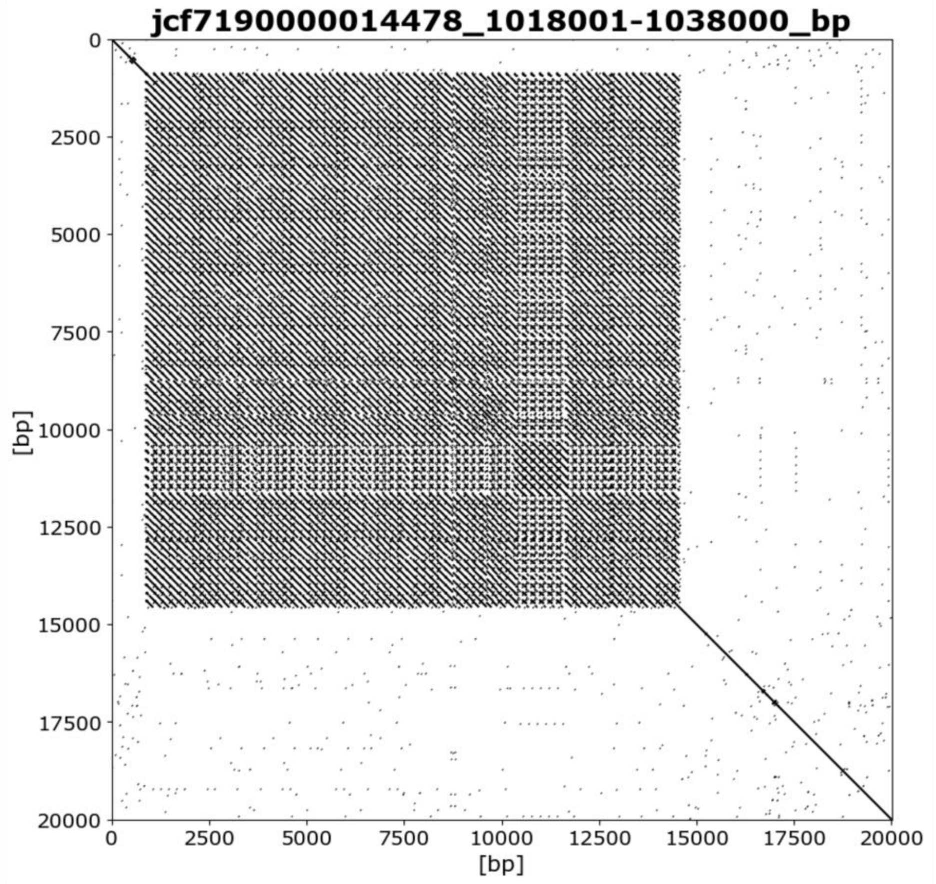


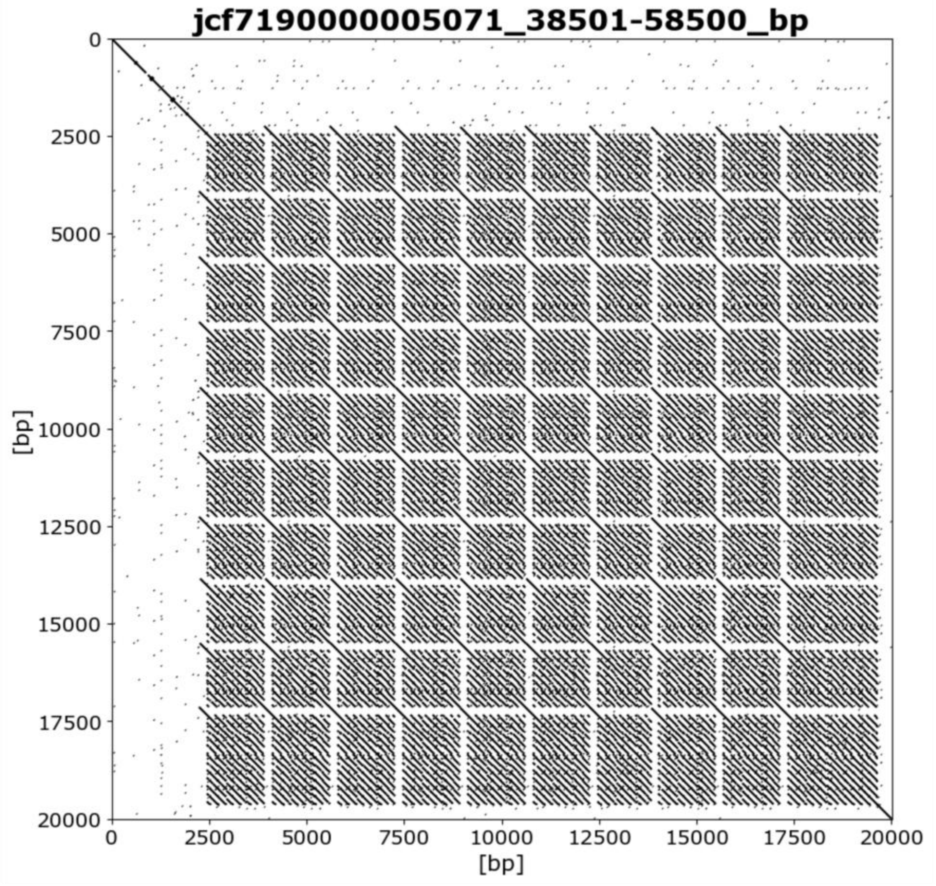

**C** EulaSat4 on *Abies alba* scaffolds (exemplary extractions of 20 kb)

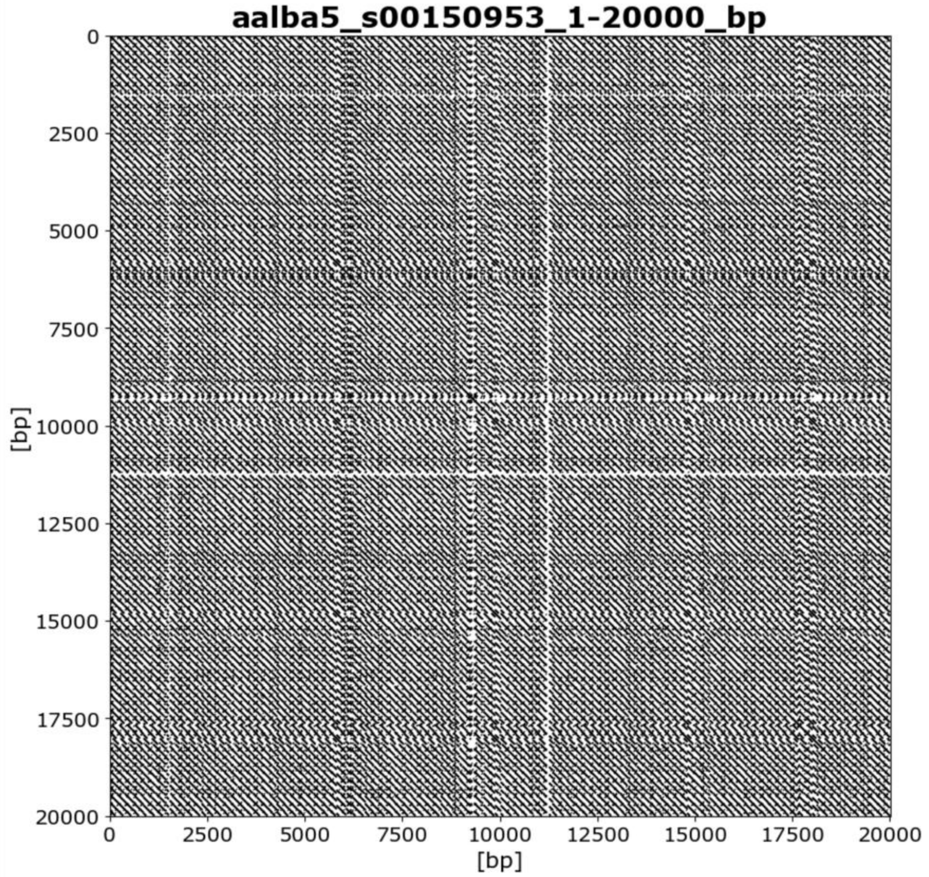


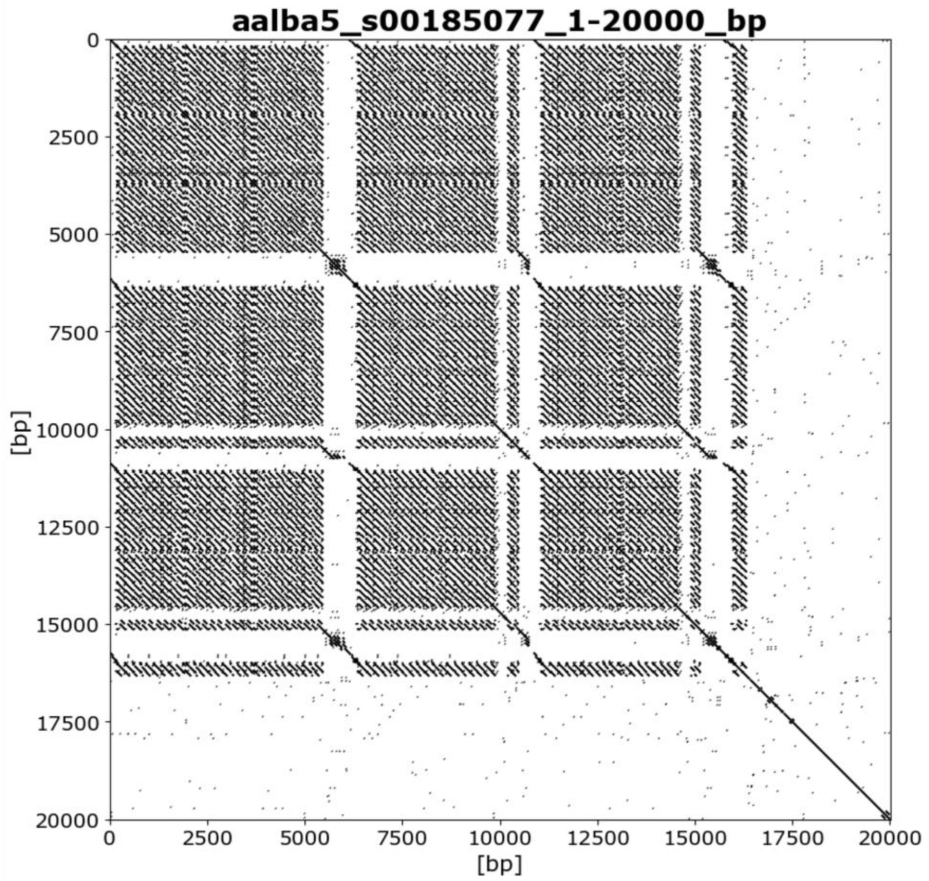


**Figure S4: Dotplots of satDNA-containing scaffolds of *P. menziesii* and *A. alba*.** Extractions of 20 kb from representative *P. menziesii* **(A, B)** and *A. alba* **(C)** scaffolds are shown, containing the EulaSat2 **(A)** and EulaSat4 **(B, C)** tandem repeats. Dotplot representations illustrate the repeat organization on scaffolds. Sequence similarities exceeding 18 bp with allowance of four mismatches are displayed as parallel lines.

**Data S1:** Consensus sequences of the EulaSat monomers in fasta format.

```
>EulaSat1_Larix_decidua
TTTCATAAATGGAATCAACAAAGTATGCACATTCTACGTCATAACGACTTTCTATAGTTTGCGCATGCGTCCGGAAATAAGAAAAGCTTA
CTTCCCCCGTTTTTTAAAATCACAGCTTCTAGAAGGTTTACATGATTTCTTAAAAAACACGAGTTTTTAGAAAATGTGTTTTA
>EulaSat1_Larix_kaempferi
TTTCATAAATGGAATCAACAAAGTATGCACATTCTACGTCATAACGACTTTCTATAGTTTGCGCATGCGTCCGGAAATAAGAAAAGCTTA
CTTCCCCCGTTTTTTAAAATCACAGCTTCTAGAAGGTTTACATGATTTCTTAAAAAACACGAGTTTTTAGAAAATGTGTTTTA
>EuLaSat2_Larix_decidua
AAAATAGCTCGGAACGTCACGAAAGTTGGCGTGGACGCTTGTCTACCAATGGGACATCCAAATCTATTCTCAAATTTCAATTCCGAGAAG
TTGGTCAAAGTTGAAACTCGACCGTGCGTTTTCGCTAGACTTGGGCTTAAAGGGTGAG
>EuLaSat2_Larix_kaempferi
AAAATAGCTCGGAACGTCACGAAAGTTGGCGTGCACGCTTGTCTACCAATGGGACATCCAAATCTATTCTCAAATTTCAATTCCGAGAAG
TTGGTCAAAGTTGAAACTCGACCGTGCGTTTTCGCTAGACTTGGGCTTAAAGGGTGAG
>EulaSat3a_Larix_decidua
CTTTTTGGATTTTTTTAGGTTACTTAAAAAAACCAAAAAAACACTTTTTTTGGCCATTTAAATCATTCAAATTACCTTTATAAGTGTATG
GCAAGATAGCTAAACGACTATGCTAACATTTCTGACCATAATTAGCATTCTAAATGCATTTTTGGAAAAATTCTGAATTTTTTAGTGTGA
TTGTTCAGTAGTTGCCACTTCAGTCAAACCATATAAACATTTAAAAATCTAACTTACATTGATCCAATCATCTTCAAACCTATATCAAAG
TCTAGCTGAACGTTTTCTGAATCTTTTAATCCAGATTTCAGCTCTCAAAGCCTTTTAGTTTATTATTTATAAGCA
>EulaSat3b_Larix_decidua
TATATGTAAATTTCTTGGGTAAGTCACAAAAATCAAAAAACAACTCAATTTGACAATAAGATCCAACCCAATCTGCATTAAATGTATATG
GCTAGTTATGTTAACATATTATCTAGACTTTTATACCATATCCAGTACTTCTAGAGCTCTTTTTTAGAAAATAGGAATTTTTTAGTGTAA
TTGTTCAGTAGTTGTCAACACAATTAACCAAGTAAACAACATTTTTTTCTAACTTCTACAACACCAATCAACTTGAAACCTATCTGAAAG
GCTAGCTGATGATTTTCTGAATATTTTGACACTACTCACAGTTTCCTAGGACTCATTTTGTGGAACCTATGGCCA
>EulaSat3c_Larix_decidua
TTTTAGTGAATTTTGTAGGTTATTTAAAAGAAACAAACAAACACTATTTTTTCCCATTTAATACTGTCAAAAAACTTTTATATGTGTATG
GAAAGACAGATAAACGTGTTTTCTTACTTTTCTGACCAAAATTAGCATTCTAAATGCAATTTTGGAAAAAATCTGAATTTTTTTGTGTTT
TTGTTCAGTAGTTGCCATCACAGTAAAAACATATAAACATTAAAAAATGTAAGTTATATTGATCCAATCATCTTAAAAACTATATCAACG
TCTAGCTAAACGTTTTCTGAATCTTTTGGTCCAGATTTGAGCTCTCTAAGTCTTTTTTTATACAATTTATAAGCA
>EulaSat4a_Larix_decidua
TTCAAAATAGAGCACATGGCACAAGCTCAAGGTATAAGCTAGCAACCACCAATCACCATGGACAGGGTTTTTCTATTGGAAAGCTAGCGA
CTGCTAGCTTTCCTTCTAGCATAAGCATGTGTGTTTATGCTAGAAGGAAATGCTTATGCTAGAAGGAAAGCTAGCAGTCGCTAGCTTTCC
TTGATCTTGTGCCATGTGCTTAT
>EulaSat4a_Larix_kaempferi
TTCAAAATAGAGCACATGGCACAAGCTCAAGGTATAAGCTAGCAACCACCAATCACCATGGACAGGGTTTTTCTATTGGAAAGCTAGCGA
CTGCTAGCTTTCCTTCTAGCATAAGCATGTGTGCTTATGCTAGAAGGAAATGCTTATGCTAGAAGGAAAGCTAGCAGTCGCTAGCTTTCC
TTGATCTTGTGCCATGTGCTTAT
>EulaSat4b_Larix_decidua
TTCAAAATAGAGCACATGGCACAAGCTCAAGGTATAAGCTAGCAACCACCAATCACCATGGACAGGGTTTTTCTATTGGAAAGCTAGCGA
CTGCTAGCTTTCCTTCTAGCATAAGCATGTGTGTTATGCTAGAAGGAAATGCTTATGCTAGAAGGAAAGCTAGCATTCG
>EulaSat4b_Larix_kaempferi
TTCAAAATAGAGCACATGGCACAAGCTCAAGGTATAAGCTAGCAACCACCAATCACCATGGACAGGGTTTTTCTATTGGAAAGCTAGCGA
CTGCTAGCTTTCCTTCTAGCATAAGCATGTGTGTTATGCTAGAAGGAAATGCTTATGCTAGAAGGAAAGCTAGCATTCG
>EulaSat5_Larix_decidua
AGTCCAGGGATGATCCAATCCCCTCAACTGTCCACTAAGGACTTCATTCTCGGAGACCTCACGTCTACGGCTCTCTTTAGGACTGAA
>EulaSat5_Larix_kaempferi
AGTCCAGGGATGATCCAATCCCCTCAACTGTCCACTAAGGACTTCATTCTCGGAGACCTCACGTCTACGGCTCTCTTTAGGACTGAA
```
